# Genomic properties of structural variants and short tandem repeats that impact gene expression and complex traits in humans

**DOI:** 10.1101/714477

**Authors:** David Jakubosky, Matteo D’Antonio, Marc Jan Bonder, Craig Smail, Margaret K.R. Donovan, William W. Young Greenwald, Agnieszka D’Antonio-Chronowska, Hiroko Matsui, i2QTL Consortium, Oliver Stegle, Erin N. Smith, Stephen B. Montgomery, Christopher DeBoever, Kelly A. Frazer

## Abstract

Structural variants (SVs) and short tandem repeats (STRs) comprise a broad group of diverse DNA variants which vastly differ in their sizes and distributions across the genome. Here, we show that different SV classes and STRs differentially impact gene expression and complex traits. Functional differences between SV classes and STRs include their genomic locations relative to eGenes, likelihood of being associated with multiple eGenes, associated eGene types (e.g., coding, noncoding, level of evolutionary constraint), effect sizes, linkage disequilibrium with tagging single nucleotide variants used in GWAS, and likelihood of being associated with GWAS traits. We also identified a set of high-impact SVs/STRs associated with the expression of three or more eGenes via chromatin loops and showed they are highly enriched for being associated with GWAS traits. Our study provides insights into the genomic properties of structural variant classes and short tandem repeats that impact gene expression and human traits.

## Introduction

Structural variants (SVs) and short tandem repeats (STRs) are important categories of genetic variation that account for the majority of base pair differences between individual genomes and are enriched for associations with gene expression (Chiang et al., 2017; Schlattl et al., 2011; Sudmant et al., 2015). SVs and STRs are comprised of several diverse classes of variants (e.g., deletions, insertions, multi-allelic copy number variants (mCNVs), and mobile element insertions (MEIs)), and multiple algorithmic approaches and deep whole genome sequencing are required to accurately identify and genotype variants in these different classes (Jakubosky et al., 2019). Due to the complexity of calling SVs and STRs, previous genetic association studies have generally not identified a comprehensive set of these variants but rather have focused on one or a few of the class types, and therefore the genomic properties of SVs and STRs associated with gene expression and/or complex traits are not well characterized.

One important problem that has not been addressed by previous SV and STR studies is whether the diverse variant classes differ in their functional impact on gene expression (DeBoever et al., 2017; Li et al., 2017; Sudmant et al., 2015; Willems et al., 2017). Therefore, while SV classes and STRs vary in genomic properties including size, distribution across the genome, and impact on nucleotide sequences, it is unknown whether these differences result in the variant classes having differential impact on gene expression such as their likelihood of being associated with a gene (eGene), their effect size, and the types of associated eGene (e.g., coding, noncoding, level of evolutionary constrant). Further, it is unknown if the variant classes exert their effects on gene expression through different mechanisms such as directly altering eGene copy number or altering three-dimensional spatial features of the genome. A comprehensive SV and STR dataset generated using high-depth whole genome sequencing (WGS) from a population sample with corresponding RNA-sequencing data could be used to compare how the different genomic properties of SV classess and STRs correspond to differential functional impacts on gene expression.

SVs and STRs have also been associated with complex traits, though they have been studied considerably less often in GWAS than single nucleotide variants (SNVs), and the overall contribution of SVs and STRs to complex traits is not well understood (Beck et al., 2015; Brandler et al., 2018; Den Dunnen, 2017; King et al., 2015; Lupski, 2015; Malhotra and Sebat, 2012; Mirkin, 2007; Nelson et al., 2013). One difficulty with studying differences between SV classes and STRs in GWAS is that it is unknown whether the variant classes are differentially tagged by SNVs on genotyping arrays. A collection of hundreds of subjects genotyped for a full range of SVs, STRs, SNVs, and indels could be used to determine whether any particular classes of SVs and STRs are not currently captured by genotyping arrays and indicate “dark” regions of the genome not assessed by array-based GWAS. A comprehensive set of variants with corresponding linkage disequilibrium (LD) estimates would also be valuable for assessing the functional impact of SVs and STRs on complex traits using existing SNV-based GWAS.

In this study, as part of the i2QTL Consortium, we used RNA-seq data from induced pluripotent stem cells (iPSCs) from the iPSCORE and HipSci collections (DeBoever et al., 2017; Kilpinen et al., 2017; Panopoulos et al., 2017a) along with a comprehensive call set of SVs and STRs from deep WGS data (Jakubosky et al., 2019) to identify variants associated with iPSC gene expression and characterize the genomic properties of these SV and STR eQTLs. We observed that SVs were more likely to act as eQTLs than SNVs when in distal regions (> 100kb from eGenes) and that duplications and mCNVs were more likely to have distal eQTLs and multiple eGenes compared to other SVs classes and STRs. eGenes for mCNV eQTLs were also less likely to be protein coding and more likely to have strong effect sizes relative to other SV classes and STRs. We examined LD of SVs and STRs with GWAS variants and found that mCNVs and duplications are poorly tagged by GWAS SNVs compared to other variant classes. 11.4% of common SVs and STRs were in strong LD with a SNV associated with at least one of 701 unique GWAS traits; and deletion, rMEI, ALU, and STR lead eQTL variants were enriched for GWAS associations establishing that these variant classes have underappreciated roles in common traits. Finally, we found a highly impactful set of SVs and STRs located near high complexity loop anchors that localize near multiple genes in three dimensional space and are enriched for being multi-gene eVariants and associated with GWAS traits. This work establishes that different classes of SVs and STRs vary in their functional properties and provides a valuable, comprehensive eQTL dataset for iPSCs.

## Results

### eQTL mapping

We performed a *cis*-eQTL analysis using RNA sequencing data from iPSCs derived from 398 donors in the iPSCORE and HipSci projects along with a comprehensive map of genetic variation (37,296 SVs, 588,189 STRs, and ~48M SNVs and indels (Table S1)) generated using deep WGS from these same donors (Jakubosky et al., 2019). These variants include several classes of SVs including biallelic duplications and deletions; multi-allelic copy number variants (mCNVs); mobile element insertions (MEIs) including LINE1, ALU, and SVA; reference mobile element insertions (rMEI); inversions; and unspecified break-ends (BNDs). We identified 16,018 robustly expressed autosomal genes and tested for cis associations between the genotypes of all common (MAF ≥ 0.05) SVs (9,313), STRs (33,608), indels (~1.86M), and SNVs (~7M) within 1 megabase of a gene body using a linear mixed model approach (Figure 1A, Table S2, Methods). We detected associations between 11,197 eGenes (FDR<5%, Methods) and 10,904 unique lead variants (lead eVariants), including 145 SVs (1.3%), 140 STRs (1.3%), 2,648 indels (24.3%), and 7,971 SNVs (73.1%, Figure 1B, Table 1). We compared our eQTLs to those discovered by GTEx and 1000 Genomes and found that the number of eGenes we identified is consistent with the expected power from using 398 samples (Figure S1–S3) (Chiang et al., 2017; Sudmant et al., 2015). While SVs and STRs accounted for only 0.1% and 0.38% of tested variants respectively in our analysis, they were highly enriched to be lead eVariants (SVs: OR=17.9, p=3.3e-91; STRs: OR=4.14, p=1.5e-24; Fisher’s exact test (FET)) and collectively formed lead associations with 3.25% of eGenes (1.73% SVs and 1.52% STRs), indicating that these variant classes have a disproportionate effect on gene expression compared to SNVs and indels.

**Figure 1.**
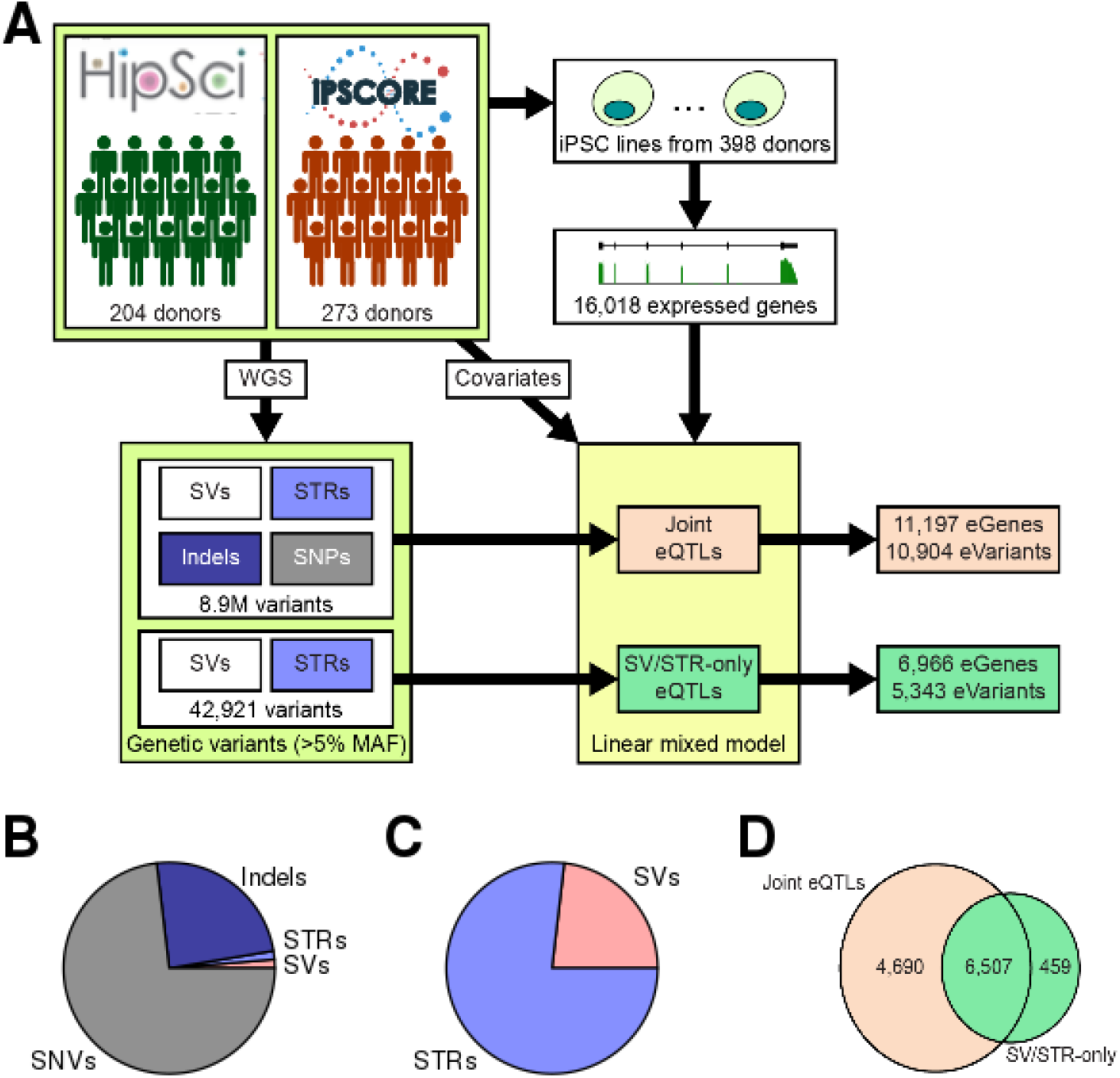
eQTL mapping. (A) Overview of eQTL study design. We used SNPs, indels, SVs and STRs called using deep WGS from 477 donors(Jakubosky et al., 2019) in conjunction with iPSCs reprogrammed from 398 of these donor(DeBoever et al., 2017; Kilpinen et al., 2017; Panopoulos et al., 2017a) and detected 16,018 expressed genes. We performed two eQTL analyses: a joint analysis that used all variants and identified 11,197 eGenes and an SV/STR-only analysis that only used SVs and STRs and identified 6,996 eGenes. (B,C) Pie charts showing the number of lead variants across the different variant classes for (B) joint and (C) SV/STR-only eQTL analyses. (D) Venn diagram showing the intersection between the eGenes detected in the joint and the SV/STR-only analysis.

**Table 1.** Summary of i2QTL variants and eQTL results. Numbers in each category refer to the number of non-redundant variants that were within 1Mb of a gene and used in the eQTL analyses. Variants used for eQTL mapping had ≥ 5% minor allele frequency for SNVs and indels and ≥ 5% non-mode allele frequency for SVs and STRs. “Lead SV/STR-Only QTLs” column shows the number of lead variants in the eQTL analysis using only SVs and STRs while “Lead Joint QTLs” column shows the number of lead variants in the eQTL analysis using SNVs, indels, SVs, and STRS.

|  | Variant Class | No. Variants | No. Common Variants | Lead SV/STR-Only QTLs | Lead Joint QTLs |
| --- | --- | --- | --- | --- | --- |
|  | SNV | 41,826,418 | 7,013,178 |  | 8,148 |
|  | INDEL | 7,040,457 | 1,862,365 |  | 2,685 |
| Copy Number Variants (CNV) | Deletion (DEL) | 16,238 | 3,073 | 661 | 51 |
|  | Duplication (DUP) | 2,693 | 391 | 55 | 9 |
|  | Multiallelic CNV (mCNV) | 1,703 | 947 | 294 | 111 |
|  | Other SV (BND) | 4,612 | 1,146 | 89 | 8 |
|  | Inversion (INV) | 210 | 84 | 11 | 0 |
|  | Reference Mobile Element Insertions (rMEI) | 2,343 | 1,448 | 243 | 3 |
| Mobile Element Insertions (MEI) | ALU | 7,880 | 1,932 | 294 | 9 |
|  | LINE1 | 1,175 | 196 | 31 | 1 |
|  | SVA | 442 | 96 | 28 | 2 |
|  | Short Tandem Repeats (STR) | 588,189 | 33,608 | 5,260 | 170 |
|  | **Total SV** | **37,296** | **9,313** | **1,706** | **194** |
|  | **Total SV/STR** | **625,485** | **42,921** | **6,966** | **364** |
|  | **Total** | **49,492,360** | **8,918,464** | **6,966** | **11,197** |

To conduct comparative analyses of the functional properties of the different SV classes and STRs, we performed an SV/STR-only eQTL analysis using the 42,921 common SVs and STRs and excluding SNVs and small indels (Figure 1A, Table S3). We identified 6,966 eGenes (FDR<5%) associated with 5,343 unique lead eVariants (Table 1, Figure 1C). SVs were more likely to be lead variants compared to STRs (OR=1.15, p=1.7e-5, FET) though the majority of lead eVariants were STRs (4,087 eSTRs versus 1,231 eSVs). Of the 11,197 eGenes identified in the joint eQTL analysis, 6,507 were also identified in the SV/STR-only eQTL analysis (Figure 1D). Among these 6,507 shared eGenes, 94.6% were mapped to a lead SNV or indel variant in the joint analysis, while the remaining 5.4% were mapped to the same lead SV or STR identified in the SV/STR eQTL analysis. To evaluate how many of the 6,155 shared eGenes were likely driven by the same causal variant in both analyses, we computed the linkage disequilibrium (LD) between SNV/indel lead variants in the joint eQTL analysis and eSVs and eSTRs from the SV/STR-only eQTL analysis. We found that lead SNVs or indels from the joint analysis were in strong LD (R^2^ > 0.8) with the lead eSV or eSTR from the SV/STR-only analysis for 14.2% (872/6,155) of shared eGenes. While the true causal variant at these loci is unknown, these data suggest that a substantial number of eQTLs that can be identified using SNVs may be explained by SVs or STRs.

### Variant size influences eQTL associations

Given that SVs and STRs have size ranges that span orders of magnitude (Jakubosky et al., 2019), we sought to examine the relationship between variant length and the likelihood of being an eVariant across the different variant classes. We tested whether STRs or deletions, duplications, and mCNVs longer than a particular length threshold were more likely to be eVariants compared to variants shorter than the length threshold. We found that longer deletions, duplications and STRs were more likely to be eVariants and lead eVariants than shorter variants (Figure 2A, Figure S4). The trend was especially strong for deletions where 38% of variants longer than 50kb were lead eVariants (OR=3.11, p=0.0065, FET). Although a higher proportion of mCNVs were eVariants compared to other classes (Figure 2A), mCNV length was not strongly associated with eQTL status; only mCNVs longer than 10kb were significantly more likely to be lead eVariants (OR=1.59, p=0.01, FET).

**Figure 2.**
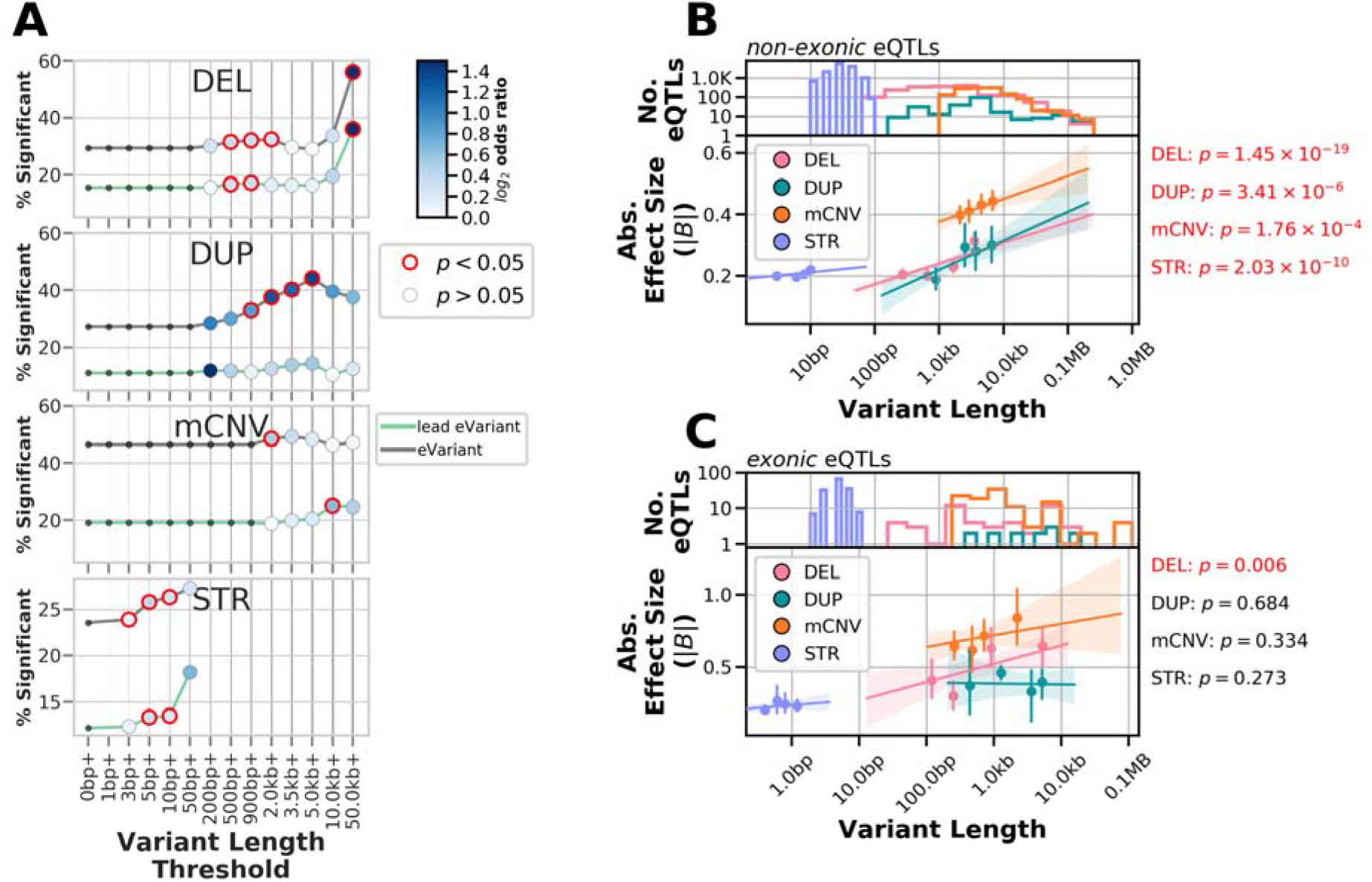
Variant length influences the likelihood and effect size of eQTLs. (A) The percentage of tested variants that were eVariants (grey lines) or lead eVariants (green lines) greater than or equal to threshold given on the x-axis for each variant class. Points are colored according to their log2 odds ratio for enrichment when comparing the fraction significant at or above the threshold to the fraction significant for variants smaller than the threshold; points circled in red were significant (FET, p<0.05). (B, C) Association of variant length with effect size for (B) non-exonic eQTLs or (C) exonic eQTLs mapped to biallelic deletions, duplications, multi-allelic CNVs and STRs. Number of eQTLs for each variant class at defined length is shown (top panels). Points represent the centers of bins with equal numbers of observations and error bars indicate 95% confidence intervals around the mean (1000 bootstraps) (bottom panels). Lines represent linear regressions, with 95% confidence intervals shaded, as calculated on unbinned data. P-values at the right of each plot indicate the significance of the association between length and absolute effect size (linear regression) when also including non-mode allele frequency and distance to TSS as covariates.

We next sought to examine whether eVariant length for SVs and STRs was predictive of absolute eQTL effect size and if lead eQTLs that overlap (exonic) or do not overlap (non-exonic) exons of the eGene displayed similar effects (Figure 2B,C). We found that lead eVariant length was significantly associated with the absolute effect size for non-exonic deletion, duplication, mCNV, and STR eQTLs independent of variant distance to the transcription start site and allele frequency (Figure 2B). However, among exonic eQTLs, only those mapping to deletions had a significant correlation between length and effect size with longer deletions having larger effect sizes (Figure 2C). These data show that longer variants are more likely to be eVariants for both SVs (excluding mCNVs) and STRs and that among eVariants that do not overlap exons, longer variants tend to have stronger effects on expression.

### mCNVs and deletions are enriched for associations with multiple eGenes

We next investigated whether SVs from particular classes were more likely to be eVariants or associated with multiple eGenes compared to STRs which comprised 78% of all tested variants and 70% of eQTLs (Figure 1C). We found that both mCNVs and deletions were more likely to be eVariants for at least one gene relative to STRs (mCNVs: OR=2.74, q=7.9e-50, FET; deletions: OR=1.31, q = 3.9e-10, FET) and were also more likely to be lead eVariants compared to STRs (mCNVs: OR=1.68, q=2.27e-8, FET; deletions: OR=1.32, q=3.7e-7, FET) (Figure 3A, Figure S5). Conversely, BNDs were less likely to be eVariants (OR=0.7, q= 4.2e-6, FET) or lead eVariants (OR=0.44, q=1.95e-12, FET) compared to STRs (Figure 3A). We next examined how often eVariants from each variant class were associated with multiple eGenes and found that, while many of the SV classes were more likely to affect multiple eGenes compared to STRs (Figure 3B), these effects were most prominent among mCNV eVariants and deletion eVariants which affected two or more genes 61.7% (271/439) and 44.4% (401/902) of the time respectively. Moreover, 31.2% (137/439) of mCNV eVariants and 13.4% (121/902) of deletion eVariants were associated with at least 4 genes compared to only 7.6% (604/7,915) of STR eVariants (Figure 3B). These results show that mCNV and deletion eVariants are more frequently associated with the expression of multiple genes compared to STRs and other SVs.

**Figure 3.**
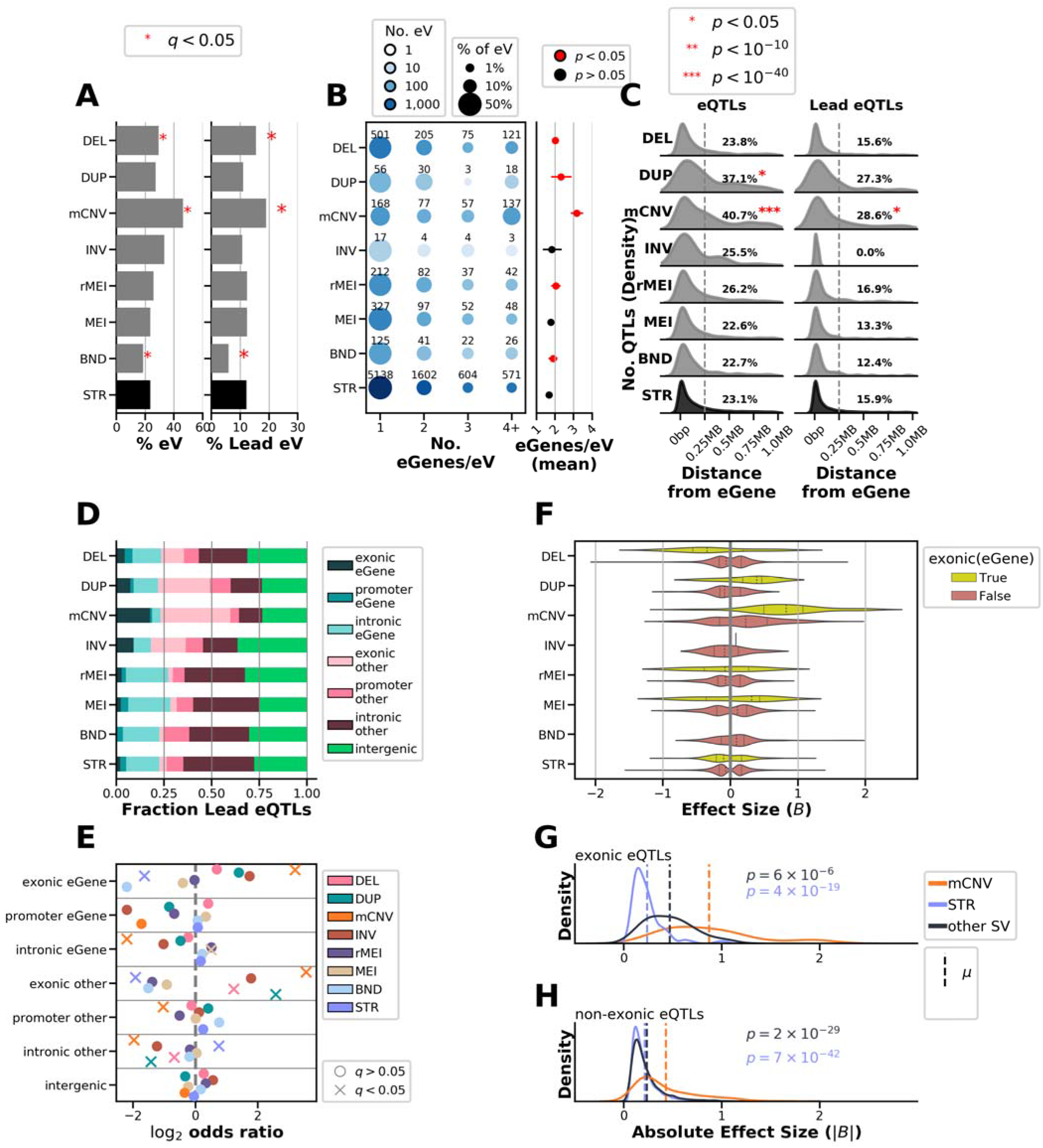
Properties of SV and STR eQTLs. (A) Percentage of tested variants from each class that are eVariants (left) or lead eVariants (right) in the SV/STR eQTL. Asterisks indicate significant enrichment or depletion, comparing the likelihood of variants from a specific class to be eVariants to that of STRs (FET, BH alpha < 0.05). (B) Balloon plot with color showing the number of eVariants and size indicating the fraction of eVariants within the variant class represented in the bin (left), and the average number of eGenes per eVariant (right) with 95% confidence intervals. Red points indicate significantly higher numbers of eGenes/eVariant (Mann Whitney U Test, Bonferonni corrected p-values < 0.05) compared to STRs. (C) Distribution of the distance of eQTL (left) and lead eQTL (right) variants to the boundary of their eGenes (5’ UTR or TSS). Proportion of eQTLs that were at least 250kb distal to eGene, red asterisks indicate that these percentages were significantly different from the other SV classes (Mann Whitney U Test, Bonferonni corrected p-values < 0.05). (D) For each variant class, the fraction of lead eQTLs that overlapped exons, promoters, or introns of their associated eGene, other genes, or that were intergenic is shown. (E) Enrichment (log2 odds ratio) comparing the proportions of lead eQTLs for each class that overlap each genic element with all other classes. (F) Distribution of effect sizes for lead eQTLs which overlapped or did not overlap an exon of their eGene. (G,H) Distribution of absolute effect sizes for mCNVs (orange), SVs excluding mCNVs (blue) and STRs (light blue) for exonic (G) and non-exonic (H) eQTLs. Annotated p-values show significantly higher effect sizes comparing the effect size distributions of mCNVs to STRs or other SVs, and are colored to match these groups (Mann Whitney U Test, Bonferonni corrected p-values < 0.05).

### Genomic localization of SV and STR eQTLs

We next examined how eVariants for each variant class were distributed with respect to genes and promoters by evaluating the distance of eVariants to their eGenes and their overlap with genic elements. We found that, for all SV classes and STRs, most eQTLs were located near eGenes (<250kb, Figure 3C); however, a significantly larger proportion of eQTLs that were mCNVs or duplications were located far from their eGenes (>250kb) compared to STRs (mCNVs: 40.7%, OR=2.23, q<1e-40; duplications: 37.1%, OR=1.93, q=4.03e-5; FET) suggesting increased distal regulatory activity for these variant types. We next annotated each variant-gene pair tested for whether the variant overlapped an exon, promoter, or intron for the paired gene; overlapped an exon, promoter, or intron for a different gene; or was intergenic (Figure 3D, Figure S6). Overall, we observed that 23.1% of lead eQTL variants directly overlapped the eGene with 205 overlapping exons (2.9%), 224 overlapping promoters (8.5%), and 1,180 overlapping only introns (17%) in the associated eGene. Interestingly, mCNVs were the only eQTL variant class whose lead variants were enriched for overlapping exonic regions of eGenes compared to all other variant classes (17.7%, OR=9.15, q=4.6e-26, Figure 3D,E). mCNV lead variants were more likely to overlap gene exons even though a substantial number of mCNV eQTLs were also located far from their eGene (Figure 3C) suggesting that a subset of mCNV eQTLs may be distal regulatory variants and a subset may affect expression by directly altering eGene copy number. Lead mCNVs, duplications, and deletions were also enriched for overlapping exonic regions of other genes besides their associated eGenes compared to other variant classes (mCNVs: OR=11.64, p<1e-40; duplications: OR=5.94,3.29e-6; deletions: OR=2.33, p=1.41e-8; FET); conversely, STRs were depleted in other gene exons (0.25%, OR=0.26, q=1.7e-16, FET, Figure 3D,E) and eGene exons (1.98%, OR=0.32, q=8.7e-14). Overall lead mCNV eVariants were more likely than other eVariants to overlap eGene exons while mCNVs and duplications had more distal eQTLs than other variant classes.

We next compared the direction and absolute effect sizes of lead eQTLs that overlapped or did not overlap exons of the eGene (exonic and non-exonic eQTLs) from each variant class to determine whether variants that alter gene copy number differ from variants that affect regulatory regions. Exonic and non-exonic lead eQTLs mapped to mCNVs and exonic lead eQTLs mapped to duplications had primarily positive associations with gene expression while exonic lead eQTLs mapped to deletions had mostly negative effects (Figure 3F). Lead eQTLs mapped to all other variant classes had bimodal effect size distributions. Comparing the absolute effect sizes of lead eQTLs mapped to each variant class, we found that mCNV lead eQTLs also had significantly larger effect sizes in both exonic (Figure 3G) and non-exonic (Figure 3H) contexts compared to lead variants from other SV classes or to STRs (Figure 3G,H). These data show that mCNV eQTLs are unique in that they tend to exert strong positive effects on gene expression, especially mCNV eQTLs that overlap exons which are almost always positively correlated with gene expression.

### eVariant type is associated with eGene type and constraint

We next investigated whether the eGene type, such as protein coding or pseudogene, was associated with the variant class of lead variants. We annotated all eGenes with Gencode gene types and calculated whether a given variant class was more or less likely to be a lead variant for eGenes of a particular gene type (Figure 4A,B, Figure S7A). Notably, a lower proportion of mCNV eGenes were protein coding (OR=0.22, q=3.33e-28, FET) and a higher proportion were pseudogenes (OR=8.56, q=1.04e-33, FET) or lincRNAs (OR=2.5, q=5e-4, FET) compared to other variant classes. Duplication and deletion eGenes followed the same trends but did not reach significance. However, STR eQTLs had the opposite pattern and were enriched for protein coding genes (OR=1.83, q=1.84e-15, FET) and depleted for pseudogenes (OR=0.36, q=2.62e-16, FET) and lincRNAs (OR=0.62, q=1.9e-3, FET). We looked at the effect sizes of associations among different gene types and found that lead eQTLs for protein coding eGenes tended to have lower effect sizes compared to lead eQTLs for genes that are not protein coding (Figure 4C) which is consistent with non-protein coding genes being more tolerant of disruption (Ruderfer et al., 2016a). Furthermore, the observation of higher effect sizes among mCNV eQTLs and their increased likelihood to overlap exons of their eGenes may be partly explained by their association with fewer protein coding genes, while the opposite properties were observed among lead eQTLs attributed to STRs which were less frequently exonic.

**Figure 4.**
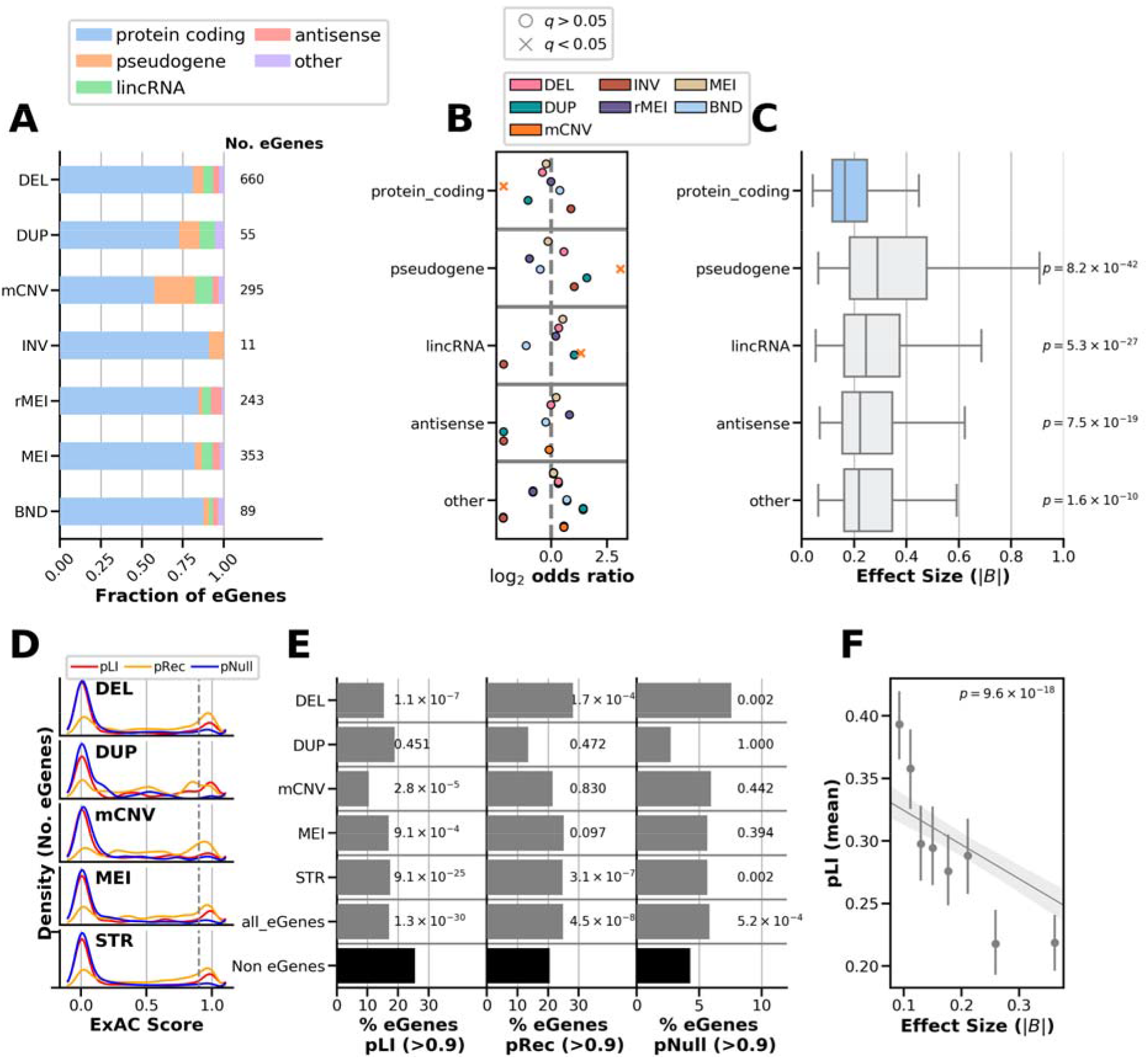
Properties of eGenes associated with different variant classes. (A) Fraction of eGenes of each Gencode subtype mapped to lead variants of each class for the SV/STR only eQTL. (B) Enrichment log2 odds ratios for the proportion of eGenes of each subtype mapped to a variant class compared with the proportion of other eGenes falling into that subtype. Significant associations (FET, BH FDR < 0.05) are indicated with ‘x’ symbols. (C) Absolute effect size of associations for genes of each subtype among lead eQTLs in the SV/STR only eQTL. p-values indicate significance of Mann Whitney U test for difference in the effect size distributions of each category as compared to protein coding genes (Bonferroni corrected). (D) Distribution of ExAC scores for intolerance to loss-of-function variants in a single allele (pLI, red), intolerance to loss-of-function variants in both alleles (pRec, orange), and tolerance to loss-of-function variants in both alleles (pNull, blue) for 5,675 eGenes. (E) The percentage of eGenes (grey bars) mapped to lead variants of each class that had high (> 0.9) pLI (left), pRec (center), or pNull scores (right). The percentage of non-eGenes (7,337 genes that were tested in the SV/STR eQTL but not significantly associated with an SV or STR) with high scores (black bars) is also included. Annotated p-values indicate the significance of the difference between the proportion of high score eGenes and high score non-eGenes for each group individually (FET, BH FDR < 0.05, within each probability score). (F) Absolute effect size versus pLI score for all eGenes after regressing out the effects of variant class and mean log10(TPM) expression level of the gene among expressed samples. Points represent binned effect size bins with equal number of observations per bin and error bars showing 95% confidence intervals (n=1000 bootstraps) around the mean pLI for the bin. Line represents a linear regression predicting pLI by eQTL effect size with variant class and the log10(TPM) expression level of the gene among expressed samples as covariates, Here the p-value represents the significance of the eQTL effect size term (t-test).

Given the differences in eGene types between different variant classes, we hypothesized that eGenes might be under different levels of evolutionary constraint compared to non-eGenes. To test this, we obtained pLI scores (probability that a gene is intolerant to loss of one allele), pRec scores (probability that a gene is intolerant to loss of both alleles), and pNull scores (probability that a gene is tolerant of loss of both copies of gene) from ExAC for 13,012 of the 16,018 genes that were tested for eQTLs (Karczewski et al., 2017; Ruderfer et al., 2016b). We examined the distributions of these constraint scores for eGenes with lead eVariants from each variant class and observed that eGenes were skewed towards low pLI scores (< 0.9) and pNull scores but more evenly distributed between low and high pRec scores (Figure 4D, Figure S7B). We found that across variant classes eGenes were significantly depleted for having high pLI scores (> 0.9) and generally enriched to have high pRec and pNull scores compared to non-eGenes (Figure 4E). This result demonstrates that genes that are intolerant to mutation are less frequently eGenes while genes tolerant of heterozygous or null alleles are more likely to be eGenes, consistent with SNV eQTLs (Lek et al., 2016). Examining this trend among variant classes, mCNVs had the lowest proportion of high pLI eGenes suggesting that mCNV protein coding eGenes are less constrained. Interestingly, eGenes mapped to deletions were most likely to be high pNull suggesting that, due to their severe negative effects on expression, deletion eVariants are under greater selection to affect dispensible genes. Given that some eGenes were classified as under high levels of constraint (pLI > 0.9), we sought to understand whether these genes are also sensitive to high levels of expression modulation. We compared the absolute effect size of lead QTLs to the pLI score of the eGene and found a strong and significant negative correlation between effect size and pLI (Figure 4F) consistent with a previous report that there is less variation in the expression of highly constrained genes (Lek et al., 2016). Taken together, these results suggest that while eGenes tend to be less constrained than other genes, the eGenes with mCNV or deletion lead eVariants are particularly tolerant of loss-of-function variation.

### Multi-eGene eQTLs Colocalize with Distal Chromatin Loop Anchors

Since chromatin looping has been shown to play a key role in the regulation of genes by positioning regulatory regions near gene promoters (Duggal et al., 2014; Greenwald et al., 2019; Rao et al., 2015; Schoenfelder et al., 2015), we sought to determine whether distal eVariants are located near the promoters of their eGenes in three-dimensional space via chromatin looping. We obtained chromatin loop calls from iPSC promoter capture Hi-C data (Montefiori et al., 2018a) that define promoter loops between gene promoters (promoter anchors) and distal sequences (distal anchors, Figure 5A). We observed that 13,575 of the 16,018 genes tested for eQTLs had at least one promoter loop and that 29.2% of SVs and 30% of STRs tested for eQTLs overlapped a distal anchor. Interestingly, mCNVs were significantly less likely to overlap a distal anchor than other variant classes with only 13.5% of mCNVs overlapping (OR=0.37, p<1e-25, FET) likely due to the difficulty of identifying loop anchors near segmental duplications, which frequently overlap mCNVs (Figure S8). Among the 13,575 genes that had at least one loop and were tested for SV/STR eQTLs, we identified 177,571 (31.8%) variant-gene pairs for which the variant was: 1) closer to the distal anchor than to the promoter anchor 2) at least 50kb away from the gene body 3) did not overlap the exon of the tested gene and 4) was a maximum of 200kb from a distal anchor, which we defined as “distal variant-gene pairs” (Figure 5A). Among these distal variant-gene pairs, we observed 1,598 eGenes (22% of all eGenes) with at least one eVariant located in the distal anchor of a loop to their promoter, 963 (12.4% of all eGenes) of which were mapped to a lead eVariant in the distal anchor; 82% (788/963) of these lead variants were STRs (Figure 5B,C). Within each variant class, distal variant-gene pairs that overlapped the distal anchor of a loop to the promoter of the tested gene were highly enriched to be eQTLs or lead eQTLs (OR=3.5, 5.4; p=0.022, <1e-40; FET, Figure 5D). These results indicate that many eQTLs include variants that overlap distal chromatin loop anchors and that variants that overlap distal anchors are more likely to be eQTLs and lead variants.

**Figure 5.**
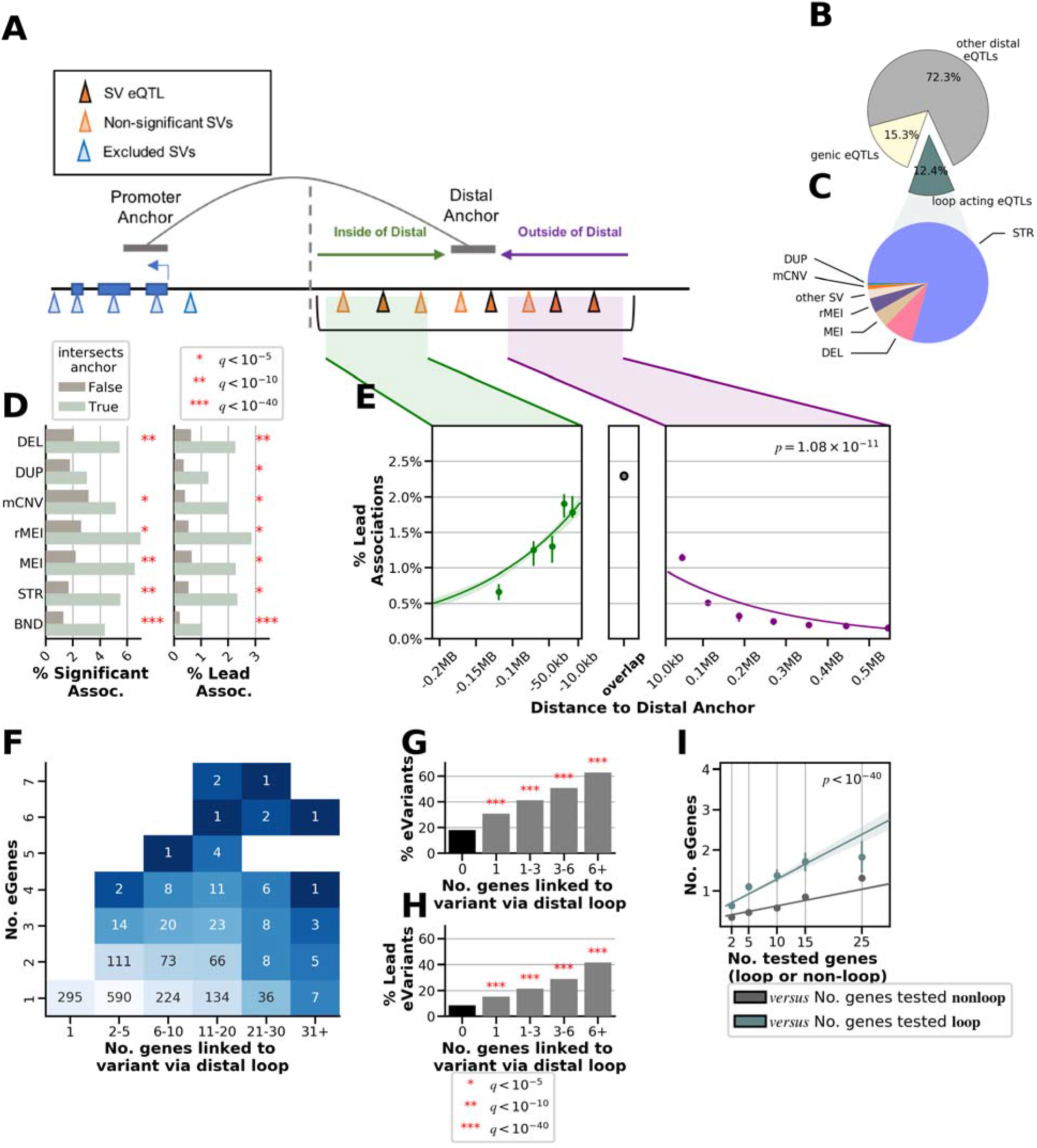
Localization of eQTLs near chromatin loops. (A) Cartoon showing localization of SVs and STRs at loop anchors. We selected all eVariants overlapping or close to distal anchors and associated with the expression of eGenes at the promoter anchor. Only eVariants closer to the distal anchor (right of grey dotted line) than to the promoter anchor were considered. (B) Proportion of eQTLs in the SV/STR eQTL analysis that were genic (overlapping an intron, exon, or promoter of the eGene; yellow), overlapping or close to distal anchors (green), or distal acting by some other mechanism (grey). (C) Distal loop-acting eQTLs (n = 2,327 eQTLs to 1,598 eGenes) mapped to SV classes. (D) Percentage of eVariant-eGene pairs where the eVariant (left) or lead eVariant (right) overlaps or does not overlap the distal anchor. Significance is calculated with FET comparing these proportions. (E) Fraction of tested distal variant-gene pairs (A) that were lead eQTLs versus their distance to the distal anchor. Points represent the centers of equally sized bins and show the mean and 95% confidence interval. Regression lines were calculated using logistic regression testing association between distance to the loop anchor (with anchors padded by 10kb) and whether the variant-gene pair was a lead association. These were computed separately for variant-gene pairs inside the loop or outside the loop. (F) Heatmap showing the number of eVariants connected to gene promoters through chromatin loops (X axis) and the number of these connected genes that are associated eGenes (Y axis). (G,H) Barplot showing the percentage of tested variants that were eVariants (G) or lead eVariants (H) as a function of the number of genes the variant was linked to via overlapping a distal anchor. P-values were calculated by FET (Benjamini-Hochberg FDR < 5%) comparing the proportions of tested variants that were eVariants for variants linked to some number of genes by chromatin loops (grey) to the proportion of variants that were eVariants among variants that were not linked to any genes by chromatin loops. (I) Number of eGenes versus the number of tested genes per eVariant stratified by whether the genes are linked by loops to the eVariant (blue) or not linked by loops (grey). We fit a linear model comparing the number of eGenes/eVariant versus the number of genes tested using whether those genes were linked by loops or not as a covariate. The p value indicates the significance of the covariate of whether the genes were or were not linked by loops.

We next hypothesized that if variants near loops are regulating gene expression, the location of variants relative to the distal anchor should be related to the chance of that variant being an eQTL. We tested if the distance of a variant to the distal anchor or the variant’s position inside or outside of the loop was predictive of whether the variant is an eVariant using a logistic model (Figure 5A). For this model, we subsetted the variant-gene pairs to those whose variants were at least 100kb away from the nearest TSS and a maximum of 200kb from the nearest distal anchor to ensure we were examining interactions around the distal anchor. We observed that variants closer to the distal loop anchor were significantly more likely to be lead eQTLs (p=0.003) and that distal variant-gene pairs with a variant inside the loop were more likely to be lead eQTLs than those with variants outside the loop (Figure 5E, OR=1.5, p=2.1e-8, FET). This suggests that variants near distal loop anchors are more likely to affect expression of the looped gene and that variants that do not directly overlap the loop anchor can still affect gene expression, potentially through changes in regulatory elements or loop structure.

Given that variants overlapping distal anchors are more likely to be eQTLs, we hypothesized that variants that are looped to multiple gene promoters would affect the expression of many of their looped targets. To examine this, we tested whether the number of looped genes to an eVariant was associated with the number of eGenes for that eVariant. We observed that variants overlapping distal anchors that were connected to multiple genes via chromatin loops tended to be multi-gene eVariants (Figure 5F). We also found that the likelihood of a variant being an eQTL or lead eQTL increased significantly as the number of genes that the variant was looped to increased (Figure 5G,H). For example, 41% of variants linked to 6 or more genes by a distal loop anchor were lead eVariants as compared to only 8.5% of distal variants that were not linked to an eGene by loop anchor (OR=7.62, p<1e-40, FET). One possible explanation of these results is that variants looping to multiple genes are located in gene-dense regions and are therefore tested for more eGenes. To address this, we compared, for each variant, the number of genes that were tested and the number that were identified as eGenes, stratified by whether the genes were connected by loops or not connected by loops, and found variants tended to have more eGenes among looped genes than genes not connected by loops (Figure 5I, p<1e-5, t-test, Methods). This trend was consistent across SV classes (Figure S9). These results suggest that variants located in high complexity loop anchors are more likely to be multi-gene eQTLs than variants simply located near many genes.

### LD tagging and GWAS associations differ between variant classes

SVs and STRs are typically not assessed in GWAS, so the contribution of classes of non-SNV variation to complex traits and diseases is currently unclear. To examine the extent by which the different SV classes and STRs have been assayed by proxy in GWAS, we calculated LD between i2QTL variants and SNVs present in the UK Biobank (UKBB, ± 50kb of each SV and STR). We observed strong LD (R^2^ > 0.8) with UKBB SNVs for a large proportion of STRs (81.7%), ALU and LINE1 elements (79% and 83.7%), and deletions (71.2%), but a markedly lower proportion of duplications (29.8%) and mCNVs (27.4%) were in strong LD with a nearby variant (Figure 6A). We stratified our analysis of duplications and mCNVs by whether they overlapped a segmental duplication (SD) and found that those that overlapped SDs were less likely to be in strong LD with UKBB variants (18.9% duplications and 16.8% of mCNVs R^2^ > 0.8) than those that did not overlap SDs (33.3% duplications and 65.7% mCNVs R^2^ > 0.8, Figure S10), indicating that poor tagging for these classes may in part be due to the presence of repetetive sequences. We also found that only 59% of multi-allelic STRs with four or more alleles were well-tagged by UKBB SNVs. These results suggest that the duplications and mCNVs are generally not assayed by proxy in GWAS, especially when located in segmental duplications, suggesting that the impact of these SV classes on traits and diseases needs to be further investigated.

**Figure 6.**
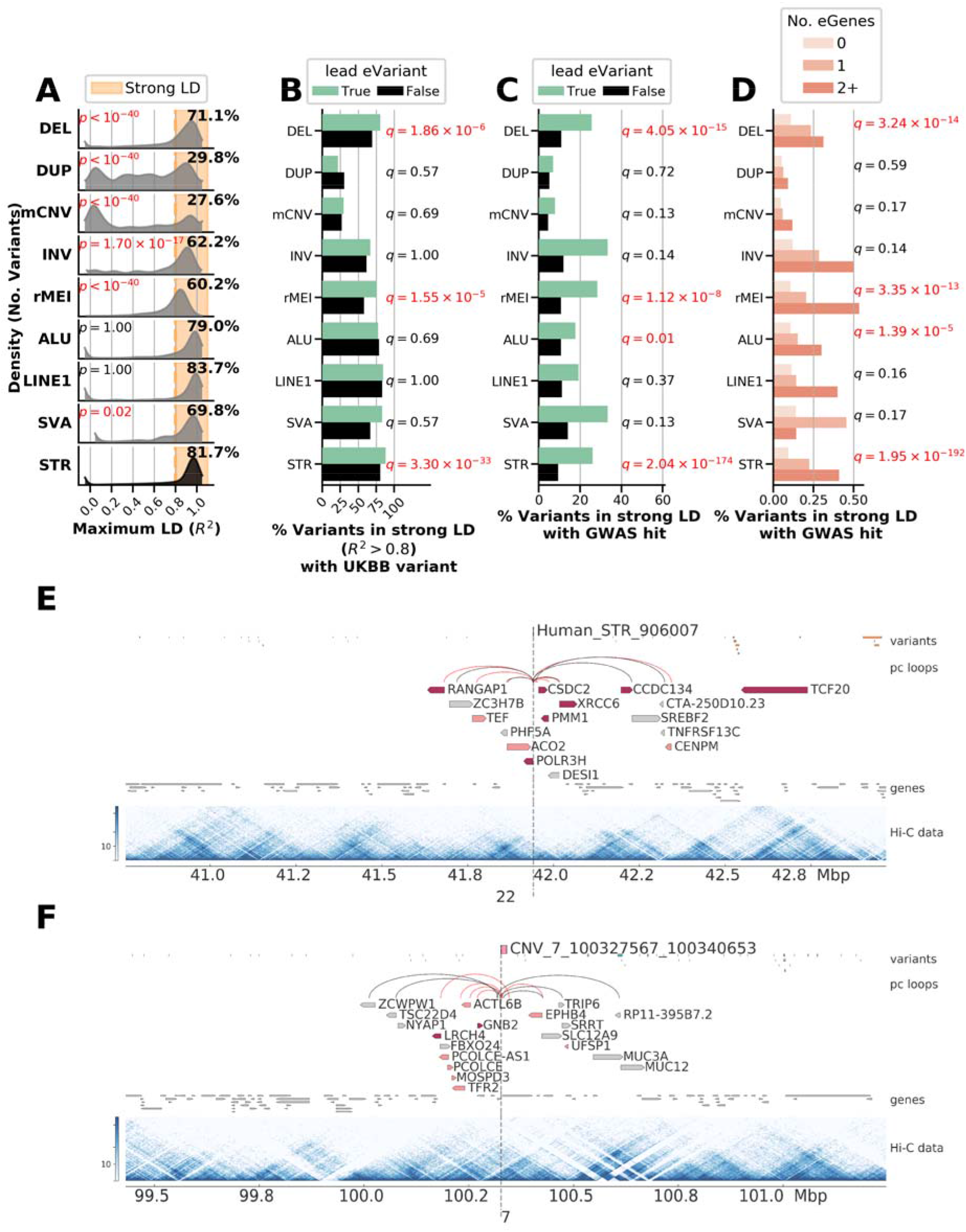
Associations between SVs, STRs and GWAS. (A) Distribution of maximum LD score per i2QTL variant with UKBB variants nearby (within 50kb) for each variant type. Annotated p-values indicate results of Mann Whitney U test for the LD distribution of each SV class to be skewed lower than STRs after Bonferroni correction for multiple tests. (B) Fraction of variants of each class that are strongly tagged by a UKBB variant (R^2^ > 0.8) for lead eVariants (green) versus all other variants in that class (black). Annotated q values indicate enrichment of lead eVariants to be in strong LD with a UKBB variant versus all other variants tested in the eQTL in the class (FET, Benjamini Hochberg FDR). (C) Fraction of variants of each class that are strongly tagged by a UKBB variant (R^2^ > 0.8) that is significantly associated with at least one trait in the UKBB (p<5e-8). Annotated q values indicate enrichment of lead eVariants to be in strong LD with a UKBB variant that is significantly associated with at least one trait versus all other variants tested in the eQTL in the class (FET, Benjamini Hochberg FDR). (D) Percentage of variants in strong LD (R^2^ > 0.8) with a variant significantly linked to at least one GWAS trait when significantly associated with 0, 1, or 2 eGenes or more. To compute annotated q-values, we utilized all variants tested in the SV/STR-only eQTL, and for each variant class we performed logistic regression to determine whether the number of eGenes for a variant was associated with whether the variant was in strong LD significant GWAS variant. The p-values from these regressions were then corrected using Benjamini Hochberg resulting in annotated q-values, marked in red when q < 0.05. (E) Example of an STR on chromosome 22 that is a multi-gene eQTL associated with a total of nine unique eGenes including four genes that the STR loops to. Genes for which the variant is a lead variant are colored dark red and genes for which the variant is significantly associated are colored pink. iPSC Hi-C data is visualized as a heatmap of interaction frequencies. The variant is located between two chromatin subdomains that span ~100kb on the left side of the variant and ~25kb on the right side of the variant (Greenwald et al., 2019). (F) Example of an mCNV on chromosome 7 that is a multi-gene eQTL associated with seven unique eGenes by looping.

Next, we investigated the extent to which SVs and STRs linked to gene expression were tagged by nearby UKBB SNVs (R^2^ > 0.8) or linked to diseases and traits via GWAS. We observed that deletions, rMEI, and STR lead eVariants were more likely to be in strong LD with UKBB variants compared to non-lead eVariants of the same class (Figure 6B, Figure S11A). While >65% of lead eVariants for most SV classes were in strong LD with any nearby UKBB variant, only 26% of mCNV and 24% of duplication lead eVariants were strongly tagged, further supporting that most mCNVs and duplications are not assayed by proxy in GWAS. We then examined how often variants in strong LD with UKBB variants were significantly associated with at least one GWAS trait (p<5e-8) and found that 11.4% of common STRs and SVs were by proxy associated with at least one of 701 UKBB traits (Figure 6C, Figure S11B). Lead eVariants were more likely to be in strong LD with significant GWAS variants, across all classes, however, enrichment was only significant in STRs, deletions, rMEIs, and ALU elements likely because other classes had too few variants to reach significance (Figure 6C). As a whole, SVs and STRs were respectively linked to 425 and 625 of 701 distinct GWAS traits, with 412 traits linked to variants of both types. Traits linked to eSVs and eSTRs were diverse including diseases such as type 1 diabetes, multiple sclerosis, arthritis, cancers, and heart disease, as well as quantitative traits such as height, body mass index and white blood cell count (Table S4).

We hypothesized that multi-eGene eVariants may have greater impact on common traits and examined the LD of these eQTLs with GWAS variants. Interestingly, we found that multi-eGene eVariants were highly enriched to be in strong LD (> 0.8 R^2^) with GWAS variants (Figure 6D). In fact, ~40% of eVariants associated with two or more eGenes were in strong LD with a GWAS variant while only 20% of eVariants associated with one eGene were in strong LD with a GWAS variant. We also observed that ~70% of eVariants associated with three or more eGenes and localized near the eGenes’ promoters via chromatin loops were in strong LD with GWAS traits. For example, a 20bp eSTR was associated with nine eGenes (seven lead) connected via distal loops (Figure 6E) and was in strong LD with a UKBB variant linked to 19 distinct traits including asthma and body fat percentage; two of the genes associated with this variant (TCF20 and POLR3H) have also been previously linked to autism (Babbs et al., 2014; Kong et al., 2012). We observed that this variant appears to overlap a chromatin subdomain boundary visible in Hi-C data from iPSCORE (Greenwald et al., 2019) which is notable given that disease causing STRs, such as the causal variant for Fragile X syndrome, have been reported to localize to subdomain boundaries (Sun et al., 2018). Additionally, we found a 13kb deletion on chromosome 7 linked to five eGenes via looping that was also linked to 14 traits (Figure 6F). These data suggest that multi-gene associations mediated by chromatin looping are frequently linked to common traits, reflecting the impact of modulating the expression of several genes.

## Discussion

We identified SVs and STRs associated with gene expression using a comprehensive SV/STR variant call set (Companion Paper) and RNA-sequencing from 398 iPSC samples. We discovered several genomic properties that were associated with gene expression across many if not all the SV classes and STRs. For deletions, duplications, and STRs, we found that increased length tended to be associated with a higher likelihood of being an eVariant and was also associated with increased effect size for non-coding eVariants. We investigated the properties of eGenes associated with SVs and STRs and showed that they were less constrained than non-eGenes and that highly constrained protein coding eGenes tended to have smaller effect sizes. Distal SV and STR eVariants were enriched for being located near the promoters of their eGenes in three-dimensional space via chromatin looping. We have previously shown that loop detection may be affected by the presence of SVs (Greenwald et al., 2019), and therefore we have likely underestimated the proportion of distal SV eVariants that mediate their effects on gene expression via chromatin loops. We also show that SV and STR eVariants near high complexity loop anchors with multiple promoter-distal regulatory element interactions are more likely to affect the expression of several genes. These results demonstrate that chromatin looping may be an important mechanism by which SVs and STRs regulate gene expression though more work is needed to establish the underlying details of this potential mechanism. Our study presents one of the largest sets of SVs and STRs associated with gene expression and reveals important general genomic properties of both SV and STR eVariants and their corresponding eGenes.

One outstanding question is whether different SV classes and STRs differentially impact gene expression. We identified substantial differences between the different SV classes in their genomic locations relative to eGenes; their likelihood of being associated with multiple eGenes; the types of associated eGene (i.e., coding, noncoding, evolutionary constraint); their effect sizes; the extent of linkage disequilibrium with tagging SNPs used in GWAS; and their likelihood of being associated with GWAS traits. Interestingly, mCNVs eQTLs differed in several respects compared to eQTLs for other variant classes. mCNVs eQTLs were more likely to be associated with the expression of multiple genes, had larger effect sizes, tended to affect noncoding genes, and were more likely to overlap the corresponding eGene or be located far from the eGene. mCNV eQTLS that overlapped exons were also highly enriched for positive associations between copy number and expression relative to other variant classes. Unlike other SV classes, the length of mCNVs was not strongly associated with the probability of being an eQTL. The differences in likelihood of being an eQTL, location, effect size, and types of eGenes for mCNVs are likely related; for instance, less constrained genes tend to have larger eQTL effect sizes, mCNVs tend to be eQTLs for less constrained genes, and mCNV eQTLs tend to have larger effect sizes. Our results indicate that a previous finding that mCNVs were enriched among predicted causal eQTL variants might be driven by the fact that mCNVs often overlap genes which likely causes differences in gene expression (Chiang et al., 2017). We also observed that deletion eQTLs were more likely to be associated with the expression of multiple genes but tended to have smaller effects on gene expression, not overlap genes, and affect less constrained genes. These observations are consistent with gene deletions and subsequent loss of expression having strong deleterious effects. Future studies may focus on whether the differences in eQTLs between variant classes are driven by selective pressures, genomic property differences between the SV classes, or some combination thereof.

The extent to which SVs and STRs contribute to variation in complex traits is not fully known because prior GWAS have generally not assessed SVs and STRs. We used our comprehensive SV/STR call set to estimate how well these variants are tagged by GWAS SNPs and whether they are associated with 701 traits from the UK Biobank. We found that only 26% of mCNV and 24% of duplication lead eVariants were tagged (R^2^ > 0.8) by a SNP in the UK Biobank, likely due in part to these variants being located in or near segmental duplications, indicating that these variants are generally missed in GWAS studies based on genotyping arrays. Multiallelic STRs are also not tagged well by SNPs and are likely not well-studied by current GWAS. We observed that 11.4% of common SVs and STRs are in strong LD with at least one significant GWAS SNP in the UK Biobank and that lead eSVs were more likely to be associated with traits compared to non-lead eSVs. We also identified a set of high-impact SVs and STRs associated with the expression of multiple genes and localized near the promoters of these genes via chromatin loops which are also highly enriched for GWAS associations. These high-impact variants that are associated with several seemingly unrelated GWAS traits may underly some of the observed pleiotropy in contempary genetic studies (Solovieff et al., 2013) and indicate that future fine-mapping efforts will greatly benefit from including SVs and STRs.

Our study demonstrates that SVs and STRs play an important role in the regulation of gene expression and that eQTLs for different classes of SVs and STRs differ in their effect sizes, genomic locations, and the types of eGenes they impact. We have also demonstrated that high-impact SVs and STRs, i.e., those associated with the expression of multiple genes via chromatin looping, are associated with a wide range of human traits. The collection of eQTLs identified here, along with the catalog of high-quality SVs and STRs described in a companion paper (Jakubosky et al., 2019), provide a powerful resource for future studies examining how these variants regulate gene expression and contribute to variation in complex traits.

## Acknowledgments

This work was supported in part by supported by the National Science Foundation, a CIRM grant GC1R-06673 and NIH grants HG008118, HL107442, DK105541 and DK112155. D.A.J. and M.K.R.D. were supported by the National Library Of Medicine of the National Institutes of Health under Award Number T15LM011271. W.W.YG. was supported by the National Heart, Lung, And Blood Institute of the National Institutes of Health under Award Number F31HL142151. S.B.M. was supported by NIH grant U01HG009431.

## Author Contributions

Conceptualization, D.A.J, E.N.S, M.D., M.J.B, S.B.M, C.D and K.A.F.; Methodology, D.A.J, M.D., M.J.B., C.S, M.K.R.D., W.W.Y.G., S.B.M, O.S., S.B.M., C.D.; Formal Analysis, D.A.J., M.D., M.J.B.; Data Curation, D.A.J., M.J.B., C.S., A.C.D., H.M.; Writing – Original Draft, D.A.J, M.D., C.D. and K.A.F.; Visualization, D.A.J; Supervision, O.S., S.B.M and K.A.F.; Funding Acquisition, O.S., S.B.M and K.A.F.

## Competing Interests

The authors declare that they have no conflicts of interest.

## Methods

### Abbreviations

1KGP: 1000 Genomes Project
eQTL: Expression quantitative trait locus. Defined by a (eGene – eVariant pair)
eGene: Gene implicated in an eQTL
eVarient: Variant implicated in an eQTL

eSV: Structural eVariant
eSTR: Short tandem repeat eVariant
eSNV: Single nucleotide eVariant
eIndel: Small insertion/deletion eVariant
FET: Fisher’s exact test
GWAS: Genome-wide association studies
Indel: Small insertion/deletion variant
LD: Linkage disequilibrium
SV: Structural Variant
SNV: Single nucleotide variant
SNP: Single nucleotide-polymorphism
WGS: Whole-genome sequencing

**Types of Genetic Variants Detected**
DEL: Biallelic deletion ascertained by LUMPY, GS, GS LCNV
DUP: Biallelic duplication ascertained by LUMPY, GS, GS LCNV
mCNV: multiallelic copy number variant ascertained by LUMPY, GS, GS LCNV. This is defined as a variant that has at least 3 predicted alleles.
INV: inversion ascertained by LUMPY
rMEI: reference mobile element insertion
BND: generic “breakend” ascertained by LUMPY. May include deletions and duplications that lack read-depth evidence, balanced rearrangements (INV), MEI or other uncategorized break points.
ALU: Non-reference Alu element insertion identified by MELT
LINE1: Non-reference LINE-1 element insertion identified by MELT
SVA: Non-reference SVA (SINE-R/VNTR/Alu) element insertion identified by MELT
STR: short tandem repeat variant, detected by HipSTR. Included variants have at least one individual with a change in length from the reference.
CNV: copy number variant (deletion or duplication structural variant). Encompasses DEL, DUP, mCNV
MEI: Non-reference mobile element insertion ascertained by MELT, including ALU, LINE1, and SVA elements

## 1. Variant Calls

Single nucleotide variant (SNV), insertion/deletion (indel), structural variant (SV) and short tandem repeat (STR) variant calls for iPSCORE and HipSci samples were discovered and rigorously analyzed in a companion paper (dbGaP: phs001325, (Jakubosky et al., 2019)).

## 2. eQTL analysis

### 2.1. RNA-Seq quality control and processing

As part of the i2QTL Consortium, we have collected a set of RNA sequencing (RNA-seq) samples from 2,954 human induced pluripotent stem cell (iPSC) lines derived from 1,600 unique donors from five studies: iPSCORE (DeBoever et al., 2017; Panopoulos et al., 2017b), HipSci(Kilpinen et al., 2017; Streeter et al., 2017), Banovich et al.(Banovich et al., 2018), GENESiPS(Carcamo-Orive et al., 2017), and PhLiPS(Pashos et al., 2017). Sample processing and quality control was performed across all samples as described below, but the eQTL analysis presented here uses a subset of the total dataset corresponding to 388 unique donors from iPSCORE and HipSci that have variant calls from deep whole genome sequencing (Jakubosky et al., 2019). The RNA-seq data were obtained from: (1) 210 iPSCORE RNA-seq samples from dbGaP (phs000924); (2) 288 HipSci cell lines (from 188 individuals) from the ENA project ERP007111 and several EGA projects (Table S5); (3) Banovich et al. (SRA: SRP093633, http://eqtl.uchicago.edu/yri_ipsc/); (4) GENESiPS (SRA – SRP072417, dbGaP: phs001139.v1.p1); (5) the PhLiPS projects (dbGaP: phs001341.v1.p1.). Data was available from these sources as either FASTQ, BAM or CRAM files. To ensure uniformity in processing, CRAM and BAM files were converted to FASTQ files. The reads in the FASTQ files were then trimmed to remove adapters and low quality bases (using Trim Galore!, http://www.bioinformatics.babraham.ac.uk/projects/trim_galore/), followed by read alignment using STAR (version: 020201) (Dobin et al., 2013) with the two-pass alignment mode and default parameters as proposed by ENCODE (c.f. STAR manual). All alignments were relative to the GRCh37 reference genome, using Ensembl 75 (Flicek et al., 2014) for any of the necessary genome annotations. Gene-level RNA expression was quantified from the STAR alignments using featureCounts (v1.6.0) (Liao et al., 2014), which was applied to the primary alignments using the “-B” and “-C” options in stranded mode when applicable. In case multiple RNA-seq runs per iPSC-line were generated these were summed to one set of gene-counts per iPSC line.

After feature quantification high quality RNA-seq samples were identified by applying filters on both Picard (https://broadinstitute.github.io/picard/) and VerifyBamID (http://csg.sph.umich.edu/kang/verifyBamID/) quality measures as well as gene expression levels. We defined high quality samples as those with > 15 million reads, > 30% coding bases, > 65% coding mRNA bases, a duplication rate lower than 75%, Median 5‘ bias below 0.4, a 3’ bias below 4, a 5’ to 3’ bias between 0.2 and 2, a median coefficient of variation of coverage of the 1000 most expressed genes below 0.8, and a free-mix value below 0.05.

Subsequently, gene expression values were normalized across lines that passed quality control. For this we derived edgeR (Nikolayeva and Robinson, 2014; Robinson et al., 2010) corrected transcript per million gene-level quantifications per iPSC line from the feature count information. After this normalization we removed samples that had low expression correlation (<0.6) with the average iPSC expression profile across our study, as measured per chromosome. This resulted in 1,378 iPSC lines derived from 1,001 donors. For the purpose of the eQTL analyses presented here, we used gene expression estimates for 288 HipSci cell lines (188 individuals) and 210 iPSCORE cell lines (210 individuals) that had corresponding deep whole genome sequencing data (WGS) that allowed for comprehensive characterization of SNVs, indels, SVs, and STRs (Jakubosky et al., 2019). This joint data set of variant calls and iPSC gene expression data for 398 individuals is referred to as the i2QTL data set in this manuscript.

### 2.2. eQTL analysis

To find eQTLs we tested for associations between variants within a cis-region spanning 1MB up- and downstream of the gene body and 16,018 robustly expressed autosomal genes (expressed in >20% of samples at an average TPM > 0.5 among samples that expressed the gene) in 398 the HipSci and iPSCORE donors (Table S1). We performed association tests using a linear mixed model (LMM), accounting for population structure and sample repeat structure as random effects (using a kinship matrix estimated using PLINK (Slifer, 2018)). All models were fit using LIMIX (Lippert et al., 2014) (https://limix.readthedocs.io/).

Before QTL testing the gene expression-levels were log transformed and standardized. Significance was tested using a likelihood ratio test. To adjust for global differences in expression across samples, we included the first 50 PEER factors (calculated across all 1,378 lines using log transformed expression values) as covariates in the model. In order to adjust for multiple testing, we used an approximate permutation scheme, analogous to the approach proposed in (Ongen et al., 2016). Briefly, for each gene, we ran LIMIX on 1,000 permutations of the genotypes while keeping covariates, kinship, and expression values fixed. We then adjusted for multiple testing using this empirical null distribution. To control for multiple testing across genes, we used Storey’s q-values (Storey and Tibshirani, 2003). Genes with significant eQTLs were reported at an FDR < 5%.

### 2.3. eQTL input variants and post processing

Because there are differences in types of SVs (e.g., copy number variants, mobile element insertions) and the output of SV variant callers, genotypes were preprocessed before use in the eQTL analysis. Since some STRs are highly multi-allelic, we used the difference in the number of base pairs with respect to the reference (expansion or contraction), as computed from the sum of the “GB” format tag in the HipSTR VCF file, as genotypes for eQTL analysis (Gymrek et al., 2016). Genome STRiP CNVDiscovery and LCNVDiscovery (Handsaker et al., 2015) variants were encoded with integer diploid copy numbers (CN). SpeedSeq (Chiang et al., 2015; Layer et al., 2014) variants were encoded using their “allele balance” (AB) fractions at each genotype, which ranges from 0 to 1 based on the amount of evidence for the variant at a site, for greater sensitivity and consistency with Genome STRiP variants, which use a continuous copy number. Finally, for MEIs identified by MELT (Gardner et al., 2017), we used traditional genotypes (0/0, 0/1, 1/1) outputted by the software, as these SVs are expected to be largely bi-allelic and there is no continuous genotype outputs available. Before performing the eQTL analysis, genotypes for all SV callers (excluding MELT) were rank normalized and converted to a 0-2 scale. For MELT variants, reference, heterozygous, and homozygous alternate genotypes were converted to 0, 1, and 2 respectively. Missing genotypes from all variant callers were filled with the mean dosage among non-missing samples prior to the eQTL. GATK (McKenna et al., 2010) SNV and indel genotypes were processed in the same way as MELT variants, converting them to 0, 1 and 2.

For the eQTL analysis, we utilized 7,013,178 SNVs, 1,862,365 indels that were present at a minor allele frequency of at least 5% and 13,804 SVs and 33,608 STRs that were present at a non-mode allele frequency of at least 5% among the 398 i2QTL donors with RNA-seq, passed QC, and were within 1MB of at least one of 16,018 expressed genes (see Methods 2.2). Notably, non-mode allele frequency was used for SVs and STRs in order to account for multi-allelic variants. For STRs, the non-mode allele frequency is computed from the difference in length of a genotype from the reference, as detailed in Gymrek et al. (Gymrek et al., 2016). The structural variant call set includes variants generated from the same caller or different callers that pass QC but may be redundant (overlapping or highly correlated) (Jakubosky et al., 2019). In a companion paper (Jakubosky et al., 2019), we identified these redundant clusters and selected high quality variants to create a non-redundant set of variants; here we chose to include all variants that passed quality control filters in the eQTL analysis including those that were marked as part of a redundancy cluster in order to maximize the chances of SV associations. Additionally, STRs were required to have a 99% call rate in both iPSCORE and HipSci samples in order to be included in the eQTL to prevent batch effects from affecting eQTLs (Jakubosky et al., 2019). To compute the number of unique variants in downstream analyses, variants were annotated with the redundancy clusters they belonged to (Jakubosky et al., 2019) and variants in the same cluster were considered as a single variant. We thus tested 9,313 non-redundant SVs that were in cis windows of expressed genes. Because variants could be associated with multiple eGenes, we considered eQTLs to be an SV/eGene pair. We performed two independent eQTL analyses: 1) using STRs, SVs, small indels and SNVs (joint eQTL analysis) and 2) using only STRs and SVs (SV/STR-only eQTL analysis; Figure 1A).

### 2.4. Association of variant length with likelihood and strength of eQTLs

To test whether longer variants were more likely to be eQTLs we restricted our analysis to tested variants from each variant class that was highly polymorphic in length (spanning orders of magnitude within variant class): duplications, deletions, mCNVs, and STRs. For variants of each of these classes, we computed the fraction of variants that were eVariants or lead eVariants that were longer than a given length threshold and calculated an enrichment p-value by comparing the proportion of variants that were eVariants or lead eVariants among variants longer than the threshold to the proportion among variants smaller than the threshold (Fisher’s Exact Test, Benjamini Hochberg). To compare the association of length of variants with effect size, we utilized all significant eQTLs from each of the aforementioned variant classes and fit a logistic regression model comparing the absolute effect size of these associations with the length of the associated variant using the distance to the nearest TSS of the eGene and the non-mode allele frequency of the variant as covariates. p-values from the regression were estimated with the Wald Test and then corrected using the Bonferroni correction.

### 2.5. Properties of SV-QTLs among different variant classes

To determine which SV classes were more likely to be associated with eGenes, we compared the proportion of variants that were eVariants for a given variant class to the proportion that were eVariants for STRs (Fisher’s Exact Test, FDR < 5%, Benjamini-Hochberg). To study the localization of eSVs with respect to eGenes, we used Gencode v19 (Harrow et al., 2012) annotations to measure distance to the nearest transcription start site for tested genes, as well as categorize variants based on their overlap of introns, exons, and promoters of tested genes, or elements of other genes. A variant was considered to overlap a particular genomic if feature if it overlapped by at least one base pair and each variant was categorized hierarchically into one of the following 7 categories, in order of precedence: 1) exonic eGene, 2) promoter eGene 3) intronic eGene 4) exonic other 5) promoter other, 6) intronic other 7) intergenic (overlapping none of the features).

### 2.6. eGenes properties and constraint

Gene types were annotated using Gencode v19 data for all expressed genes. We then performed enrichment analyses comparing the proportion of eGenes of a specific type mapped to each class to the proportion among all other variant classes (Fisher’s Exact Test, Benjamin-Hochberg correction). To compare the effect sizes of associations with each gene type, we compared the distribution of effect sizes for lead associations for protein coding eGenes to those of pseudogenes, lincRNA, antisense and all other genes (Mann Whitney U Test, Benjamini-Hochberg correction). To investigate the constraint of eGenes, we obtained ExAC (v0.3.1) (Lek et al., 2016) pLI, pNull, and pRec estimates for 13,012 expressed genes and restricted our analyses to lead associations with these eGenes. We then compared the proportion of eGenes with high (> 0.9) ExAC scores mapped to either deletions, duplications, mCNV, MEI (LINE1, SVA, and Alu), STR, or all 13,012 eGenes to the proportion of genes with a high score among the 9,052 non-eGenes that were tested in our dataset using Fisher’s Exact Test and adjusting for multiple testing with the Bonferonni method. Finally, we fit a logistic model predicting the pLI of an eGene using the eGene’s absolute effect size and including variant class as a covariate to test for an association between eGene effect size and pLI.

### 2.7. eQTL localization near distal anchors of chromatin loops

To examine the localization of SVs and eSVs with respect to chromatin loops in iPSCs, we downloaded iPSC promoter capture Hi-C loop call data from a previous study (Montefiori et al., 2018b). For this analysis, we obtained loops intersecting the promoter of 13,575 out of the 16,018 expressed genes included in the eQTL analysis, of which 5,803 were eGenes. To identify variants that might affect chromatin looping, we first intersected loop calls with all annotated Gencode v19 promoters. Then, for each variant, we computed the distance from each loop anchor and retained only the variants closer to the distal anchor (i.e. the anchor that does not overlap the promoter). We subsetted this set of variant-gene pairs to those where the variant was: 1) closer to the distal anchor than the promoter anchor 2) at least 50kb from the promoter 3) at most 200kb from the distal anchor which comprised 177,571 variant-gene pairs (31.8% of all tested variant-gene pairs). For all these variants, we determined whether they were included in the Hi-C loop (i.e. between the promoter anchor and the distal anchor) or outside the loop. To test whether variants that hit distal loop anchors are enriched to be eQTLs, we categorized variants within 10kb of a loop anchor as intersecting that anchor. To calculate enrichment, we tested the proportion of eVariants that intersected a distal loop anchor to at least one expressed gene versus the proportion of eVariants that did not intersect distal loop anchors (Fisher’s exact test). Next, we used this same subset of variant-gene pairs to test whether the distance of a variant from the distal loop anchor was associated with its likelihood of being associated with gene expression. To do so, we fit a logistic model to see whether distance to the distal anchor is predictive of the likelihood of being associated with gene expression using distance to the gene body and non-mode allele frequency as covariates. We also compared the proportion of variants that were eVariants that were within 100kb from the outside of the distal loop anchor to variants that were within 100kb from inside of the distal loop anchor (Fisher’s exact test). To test whether overlapping loop anchors associated with multiple promoters was more predictive of associations with multiple eGenes than variant localization in a gene dense window, we modeled the number of eGenes versus the number of genes tested for each eVariant, stratifying by the number of genes tested that were either connected by loops to the promoter or not connected by loops to the promoter (“statsmodels.api.logit”, statsmodels v0.9.0, https://pypi.org/project/statsmodels/). To visualize these regressions, seaborn (regplot)(https://pypi.org/project/seaborn/) was used in order to divide the X axis (number of genes tested that were connected or not connected by loops) into bins with points drawn at the center of the bin showing the mean and error bars indicating 95% confidence intervals. The same bins were used for both groups in order to enable direct comparison between groups, however, each bin does not contain equal numbers of observations.

### 2.8. SV/STR tagging and GWAS associations

We downloaded summary statistics for 4,357 human traits from the UK BioBank (UKBB) GWAS Round 2 (http://www.nealelab.is/uk-biobank, August 1, 2018). For each of the 42,921 non-redundant SVs and STRs, we used bcftools(Li et al., 2009) to extract all SNPs 50 kb upstream and downstream. For each SV or STR, we calculated LD as the correlation (R^2^) with the genotypes of each surrounding SNV or indel genotyped in i2QTL WGS. We selected the variant with strongest LD overall, as well as the variant with the strongest LD that was included in the UKBB data set (if the two were different). For each UKBB variant linked to an SV or STR, we obtained p-values for the variant in all GWAS studies and considered it to be significantly associated with a trait if p < 5 × 10^−8^. For each variant type, we selected all lead SVs and STRs from the SV/STR-only eQTL analysis and tested if the lead eVariants were: 1) more likely to be in strong LD with UKBB variants in general, and 2) more likely to be in strong LD with UKBB variants significantly associated with a GWAS trait, as compared to non-lead eVariants, using the Fisher’s exact test. To test the association of multi-eGene eQTLs with the likelihood of being in strong LD with a variant significantly associated with a GWAS trait, we divided tested variants by class and modeled the likelihood of a variant being linked to a trait versus the number associated eGenes (“statsmodels.api.logit”, statsmodels v0.9.0, https://pypi.org/project/statsmodels/). p-values were calculate using the Wald test and then corrected for multiple testing using Benjamini-Hochberg FDR.

---

**Figure S1.**
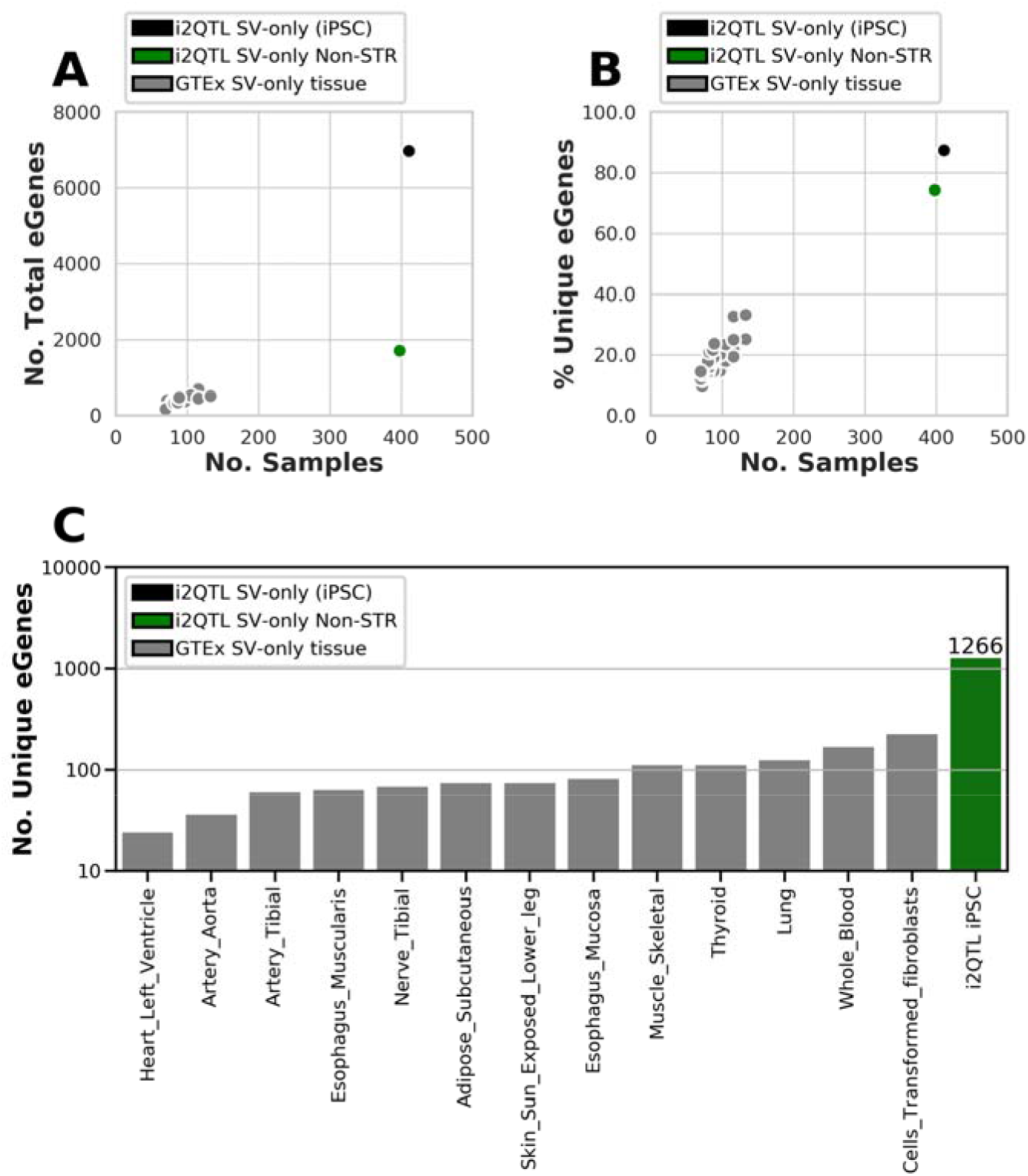
Comparison of i2QTL SV/STR-only eQTL to GTEx v6 SV only eQTL. (A) Number of eGenes and (B) percent of unique eGenes as a function of the number of samples for 13 tissues from the GTEx v6 SV-only eQTL(Chiang et al., 2017) (grey), the i2QTL SV-only eQTL (black), and the i2QTL SV-only without STRs eQTL (green). We repeated the i2QTL SV-only eQTL analysis without STRs because the GTEx study did not include STRs. (C) Number of unique eGenes for each tissue and study.

**Figure S2.**
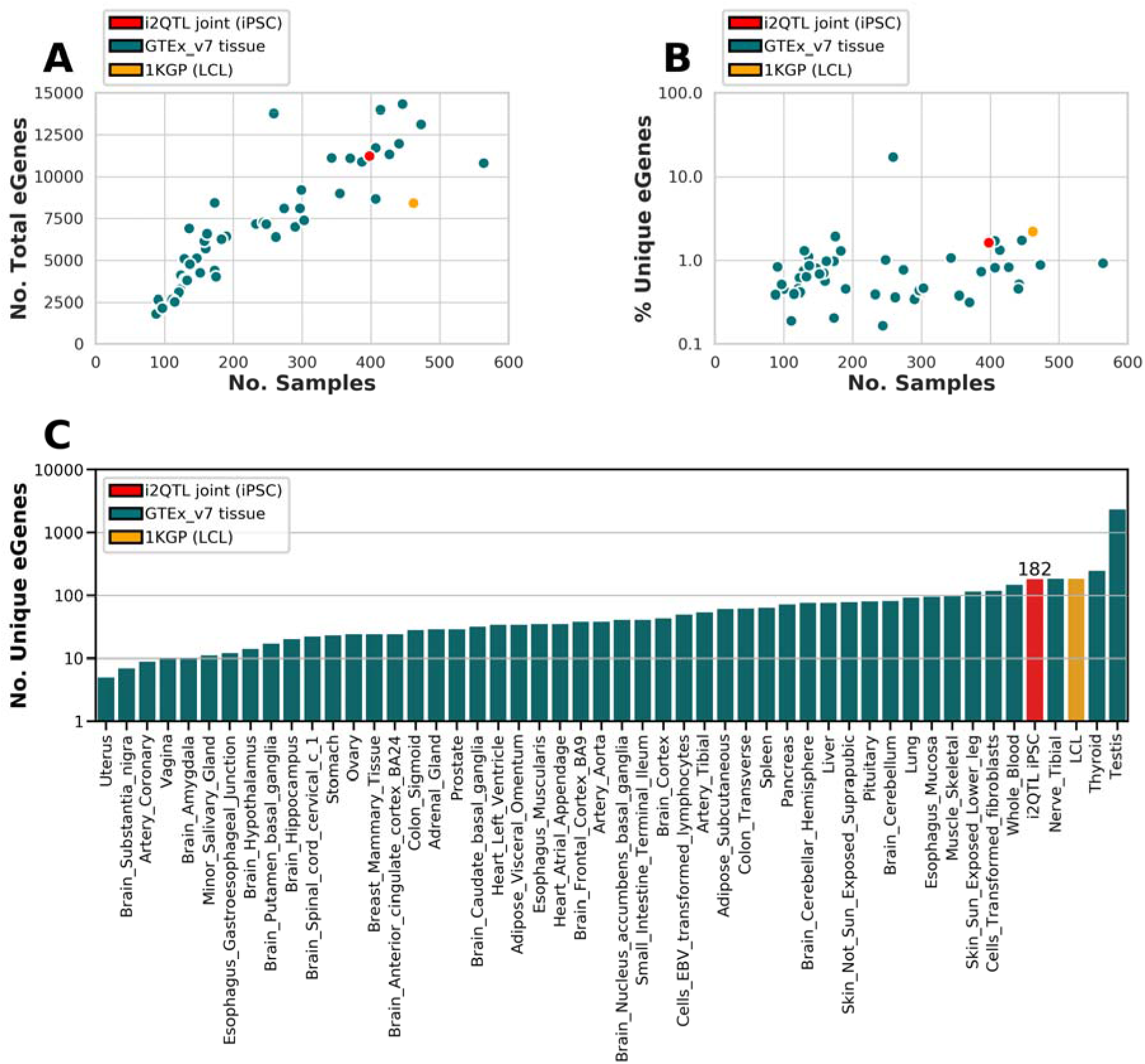
Comparison of i2QTL joint eQTL to GTEx v7 and 1KGP. (A) Number of eGenes and (B) percent of unique eGenes as a function of the number of samples for 48 tissues from the GTEx v7 eQTL (which did not include SVs, green), 1KGP SV eQTL in lymphoblastoid cell lines (orange), and the i2QTL joint eQTL that included SNVs, indels, SVs, and STRs (red). (C) Number of unique eGenes for each tissue and study.

**Figure S3.**
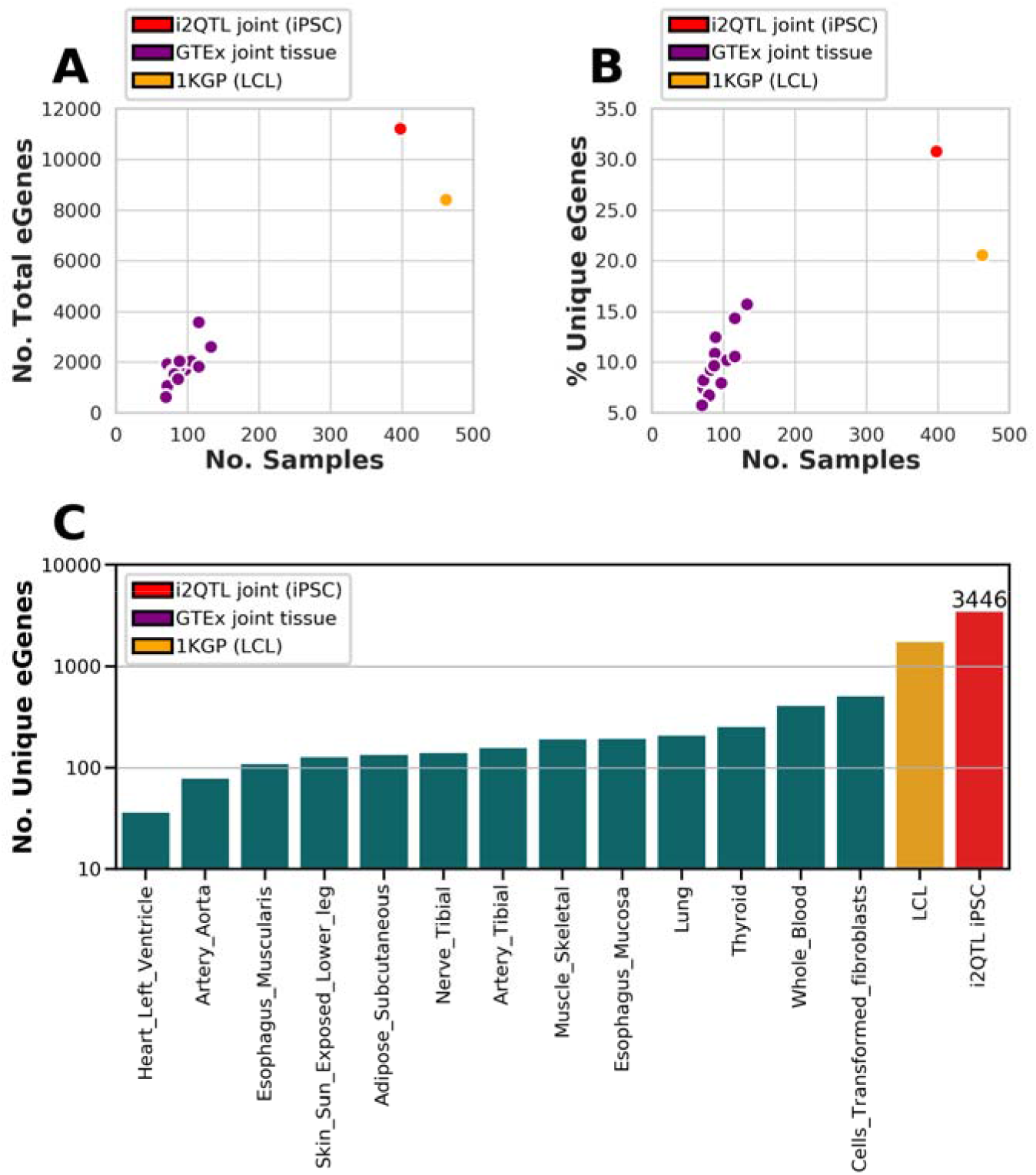
Comparison of i2QTL joint eQTL to GTEx v6 and 1KGP. (A) Number of eGenes and (B) percent of unique eGenes as a function of the number of samples for 13 tissues from the GTEx v6 joint eQTL(Chiang et al., 2017) that included SVs/SNVs/indels (puriple), 1KGP (SV/SNVs/indels)(Sudmant et al., 2015) eQTL in lymphoblastoid cell lines (orange), and the i2QTL joint eQTL that included SNVs, indels, SVs, and STRs (red). (C) Number of unique eGenes for each tissue and study.

**Figure S4.**
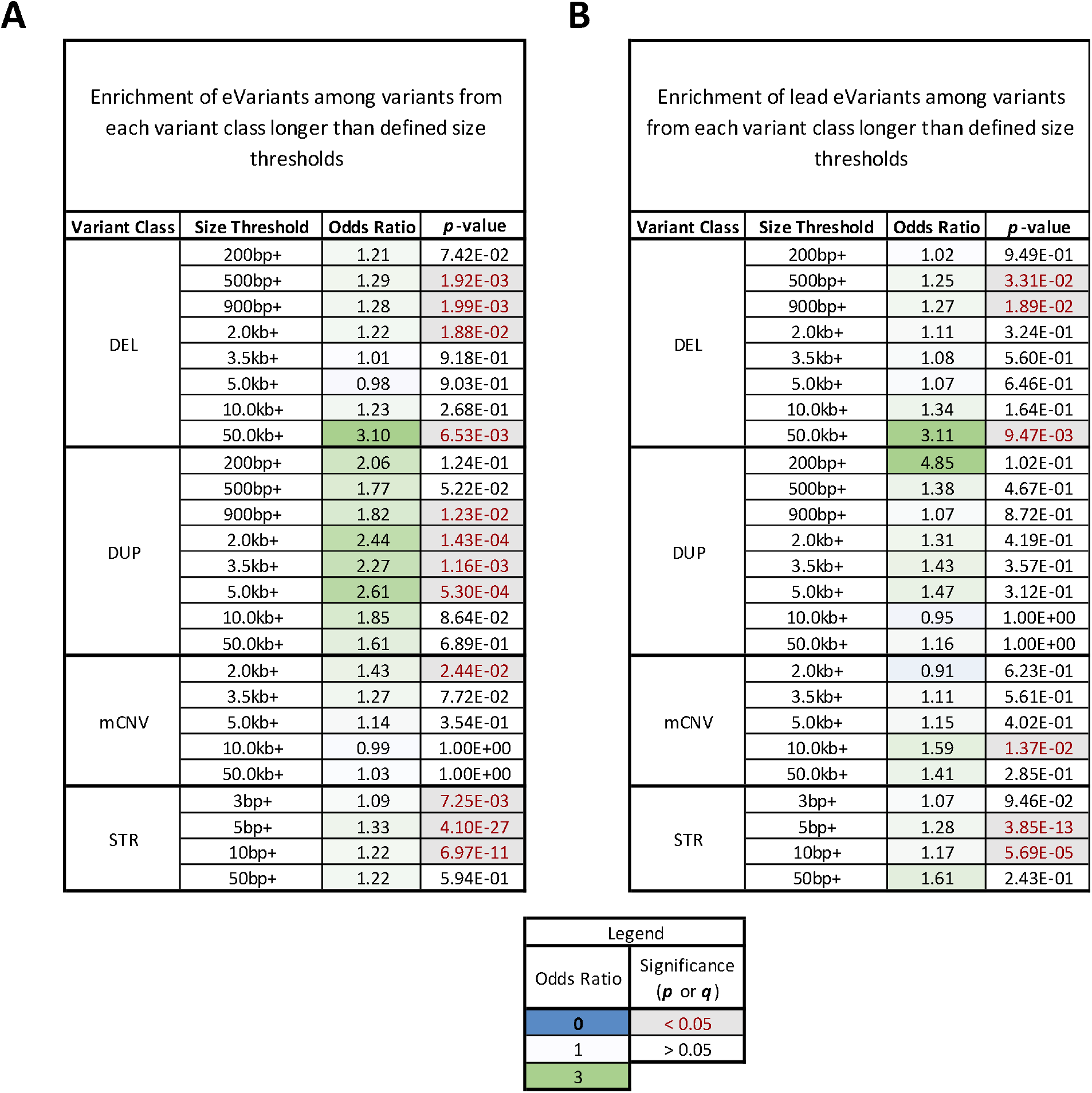
Variant Length and Likelihood of Being an eQTL. (A, B) Enrichment odds ratios and p-values for the likelihood of being an eVariant (A) or lead eVariant (B) comparing variants larger than each size threshold to those below the size threshold (Fisher’s exact test).

**Figure S5.**
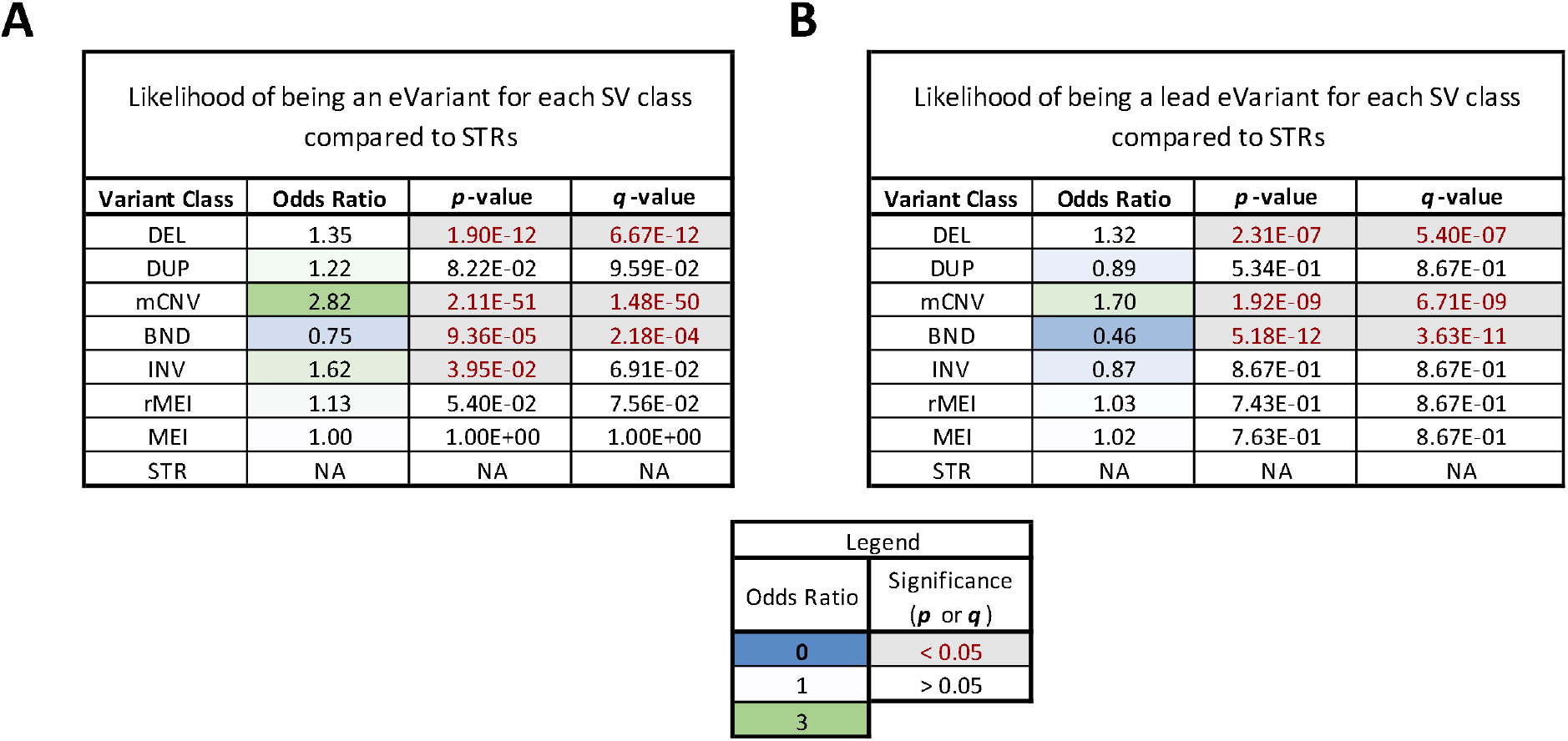
Variant Class and Likelihood of Being an eQTL. (A, B) Enrichment odds ratios, p-values, and q-values (Benjamini Hochberg) for the likelihood of variants from each class to be eVariants (A) or lead eVariants (B), as compared to STRs (Fisher’s exact test).

**Figure S6.**
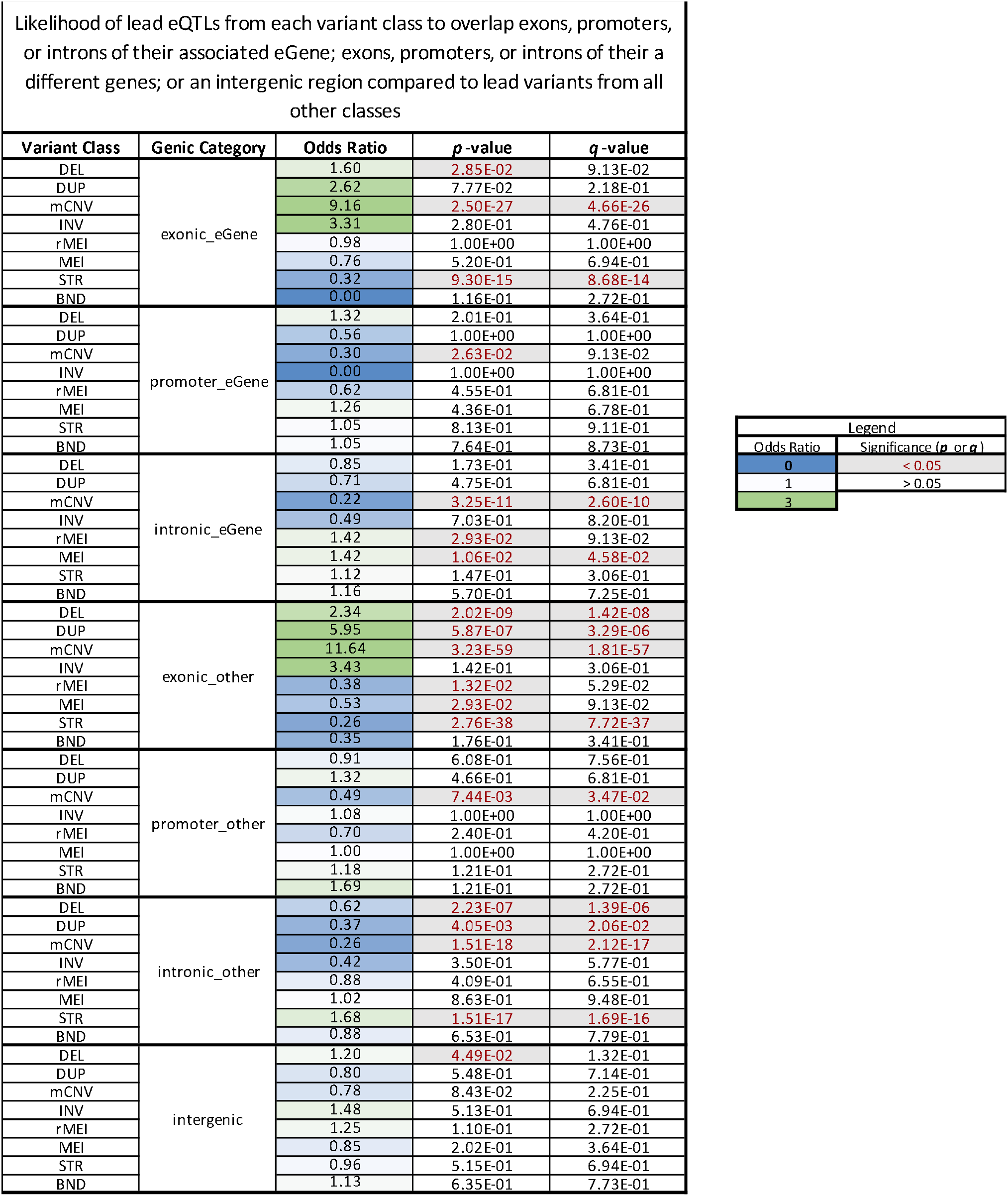
eQTL Localization with respect to genic elements. (A, B) Enrichment odds ratios, p-values, and q-values (Benjamini Hochberg) for the likelihood of eQTLs (A) or lead eQTLs (B) from each class to overlap genic elements of the eGene or some other gene as compared to all other eQTLs/lead eQTLs. Variants that overlapped a feature by a minimum of 1 base pair were assigned hierarchically in order of priority (from highest to lowest) to one of the following groups: 1) exonic to eGene 2) promoter of eGene 3) intronic to eGene 4) exonic to other gene, 5) promoter of other gene 3) intronic to other gene, and if not overlapping any of these features they were assigned as 7) intergenic.

**Figure S7.**
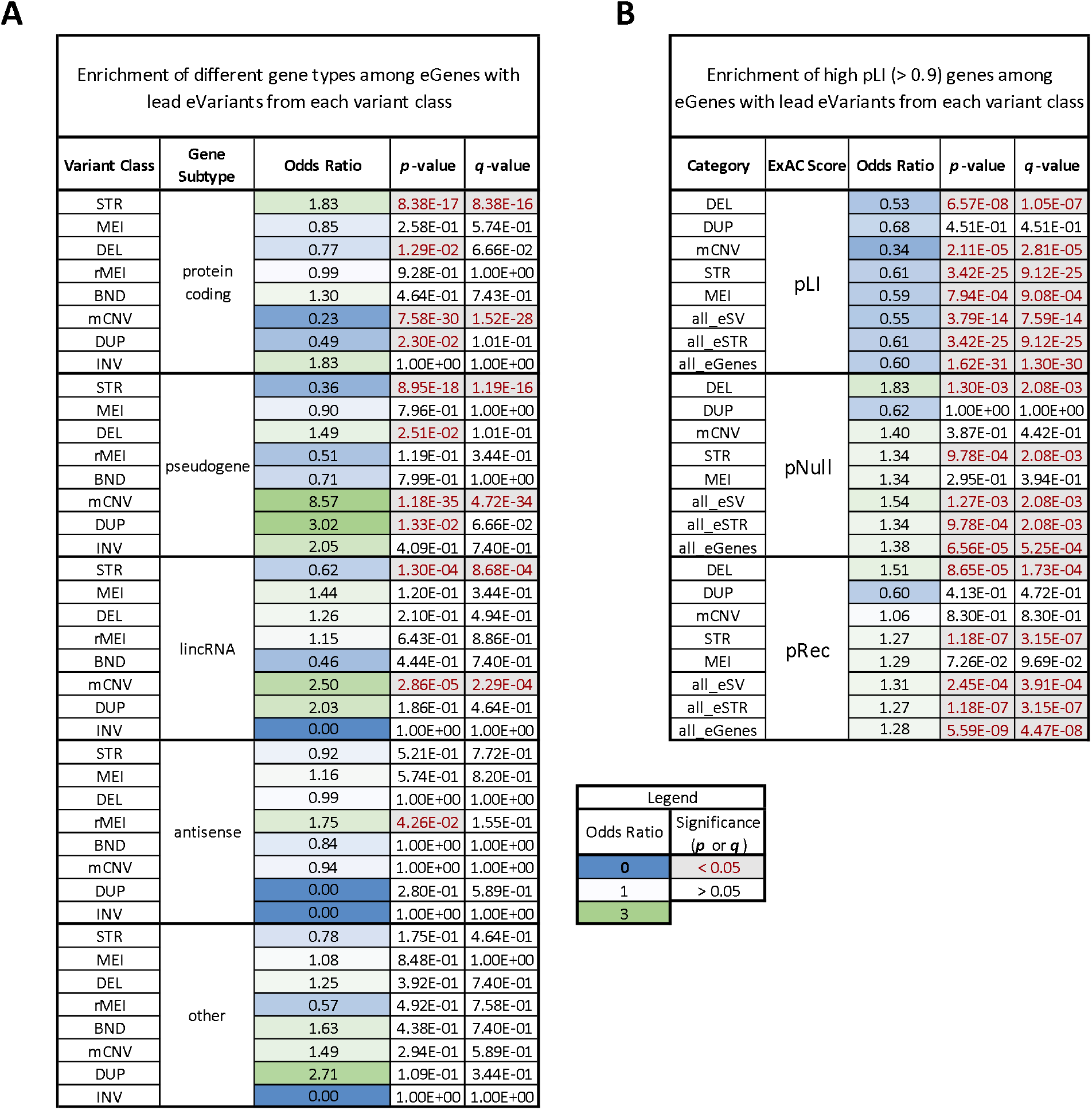
Gene type and relative constraint for eGenes mapped to different classes. (A, B) Enrichment odds ratios, p-values, and q-values (Benjamini Hochberg) for the likelihood of eQTLs (A) or lead eQTLs (B) from each class to overlap genic elements of the eGene or some other gene as compared to all other eQTLs/lead eQTLs. Variants that overlapped a feature by a minimum of 1 base pair were assigned hierarchically in order of priority (from highest to lowest) to one of the following groups: 1) exonic to eGene 2) promoter of eGene 3) intronic to eGene 4) exonic to other gene, 5) promoter of other gene 3) intronic to other gene, and if not overlapping any of these features they were assigned as 7) intergenic.

**Figure S8.**
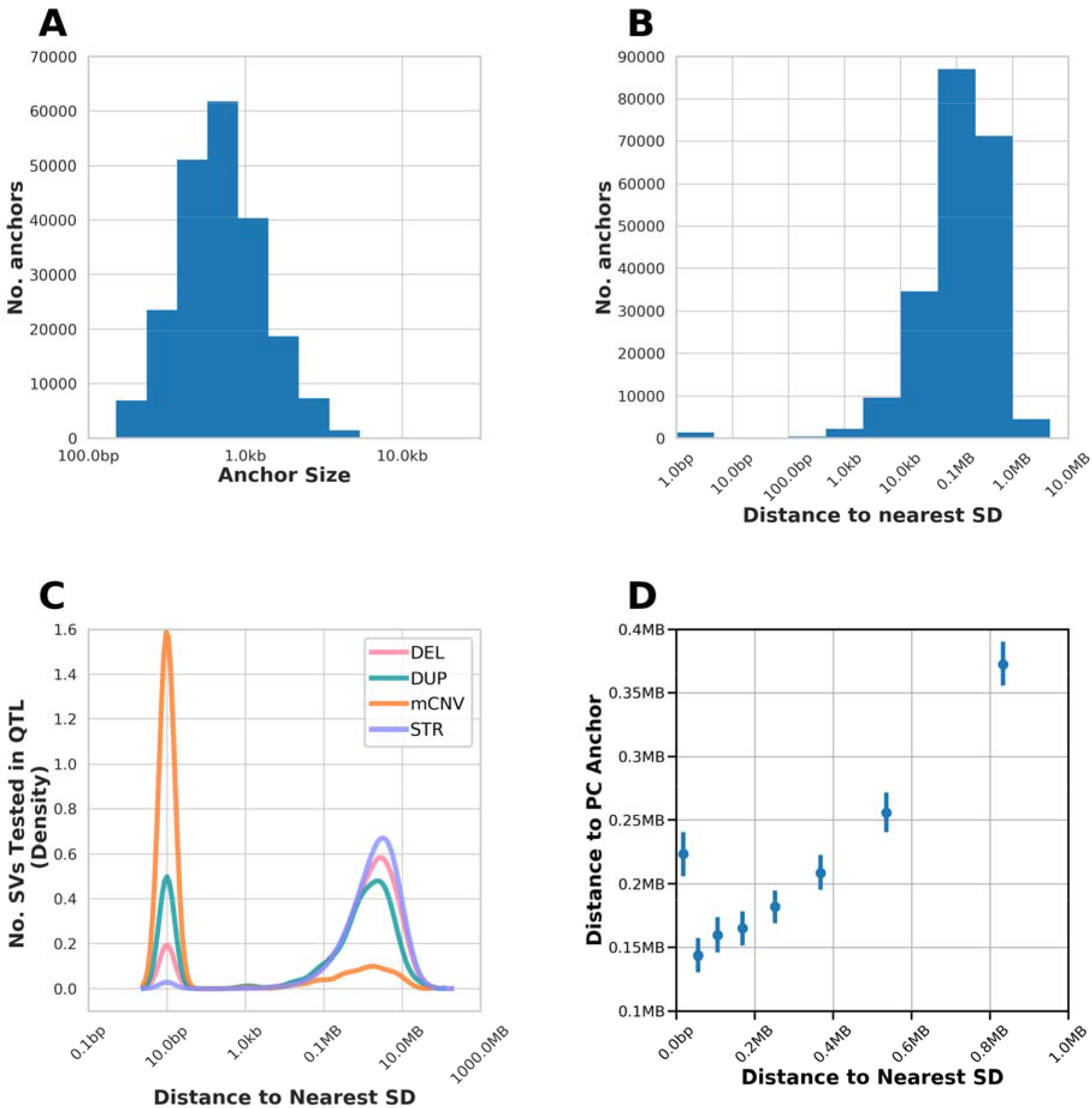
Promoter Capture Loops and Segmental Duplications. (A) Distribution of anchor sizes for unique anchors in promoter capture Hi-C data for all anchors in dataset (Montefiori et al., 2018b). (B) Distance to nearest segmental duplication (SD) for each unique anchor. (C) Distance of SVs and STRs to the nearest segmental duplication. (D) Distance to nearest segmental duplication versus distance to promoter capture Hi-C anchor for all 625,485 discovered SVs and STRs in the i2QTL call set and all Hi-C loops. Distance is binned into equally sized bins with the same number of observations per bin and error bars represent 95% confidence intervals around the mean. Variants closer to segmental duplications tend to be closer to promoter capture Hi-C anchors, however, those that overlap segmental duplications are not likely to be close to promoter capture anchors. This suggests that mCNVs that overlap chromatin loops are likely missed due to local sequence similarity within and surrounding the mCNVs precluding these loops from being identified.

**Figure S9.**
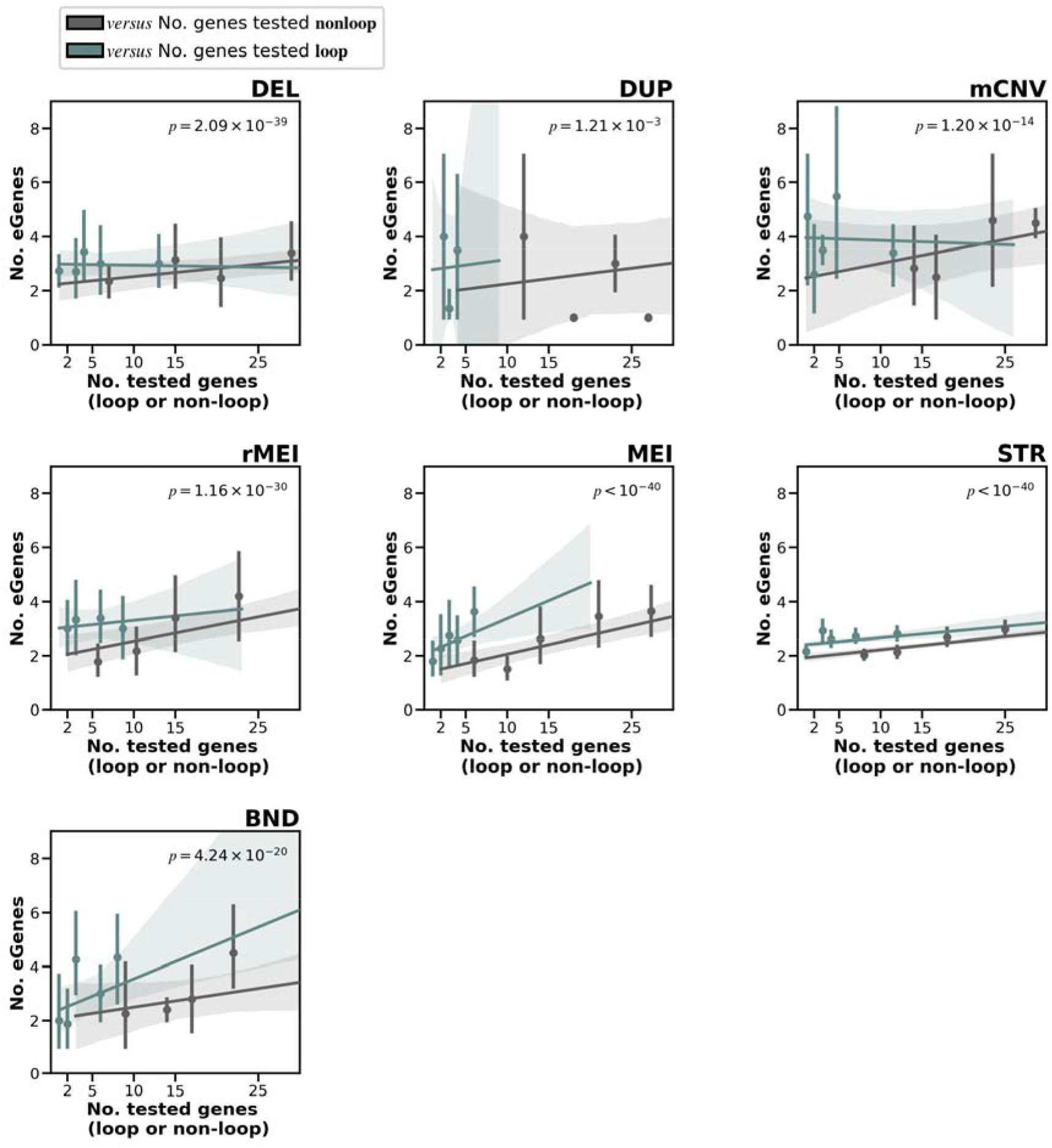
Number of eGenes versus Number of Loop Connected Genes Tested By Variant Class. Using the results of the SV/STR only eQTL, for each variant class, we compared the number of eGenes versus the number of tested genes per eVariant stratified by whether the genes are linked by loops to the eVariant (blue) or not linked by loops (grey). We used a combined linear regression model comparing the number of eGenes/eVariant versus the number of genes tested with the covariate of whether those genes were loop linked or not. The *p* value indicates the significance of the covariate of whether the genes were or were not linked by loops.

**Figure S10.**
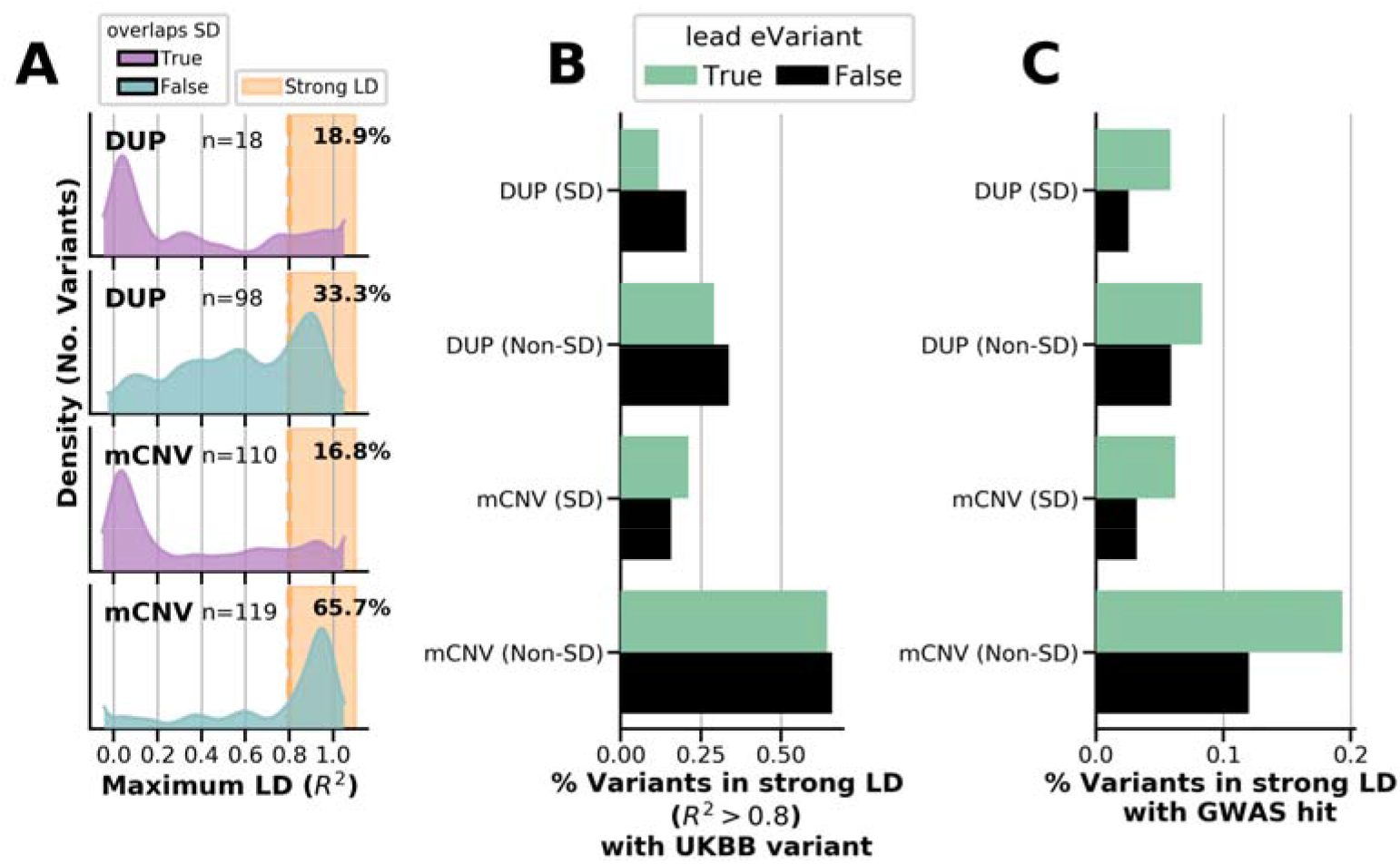
Overlap of Segmental Duplications and LD with UKBB variants. (A) Distribution of maximum LD score per i2QTL variant with nearby UKBB variants (within 50kb) for biallelic duplications and mCNVs that overlap or do not overlap a segmental duplication (SD), numbers (n=) indicate the number that are strongly tagged in each category. Annotated p-values indicate results of Mann Whitney U test for the LD distribution of the group that overlaps segmental duplications to be skewed lower than the group that does not for each variant class (B) Fraction of variants of each class that are strongly tagged by a UKBB variant (R^2^ > 0.8) for lead eVariants (green) versus all other variants in that class (black). (C) Fraction of variants of each class that are strongly tagged by a UKBB variant (R^2^ > 0.8) that is significantly associated with at least one trait (p-value < 5 × 10^−8^). Annotated q values indicate enrichment of lead eVariants to be linked to GWAS traits versus all other variants in the class (Fisher’s exact test, Benjamini-Hochberg FDR).

**Figure S11.**
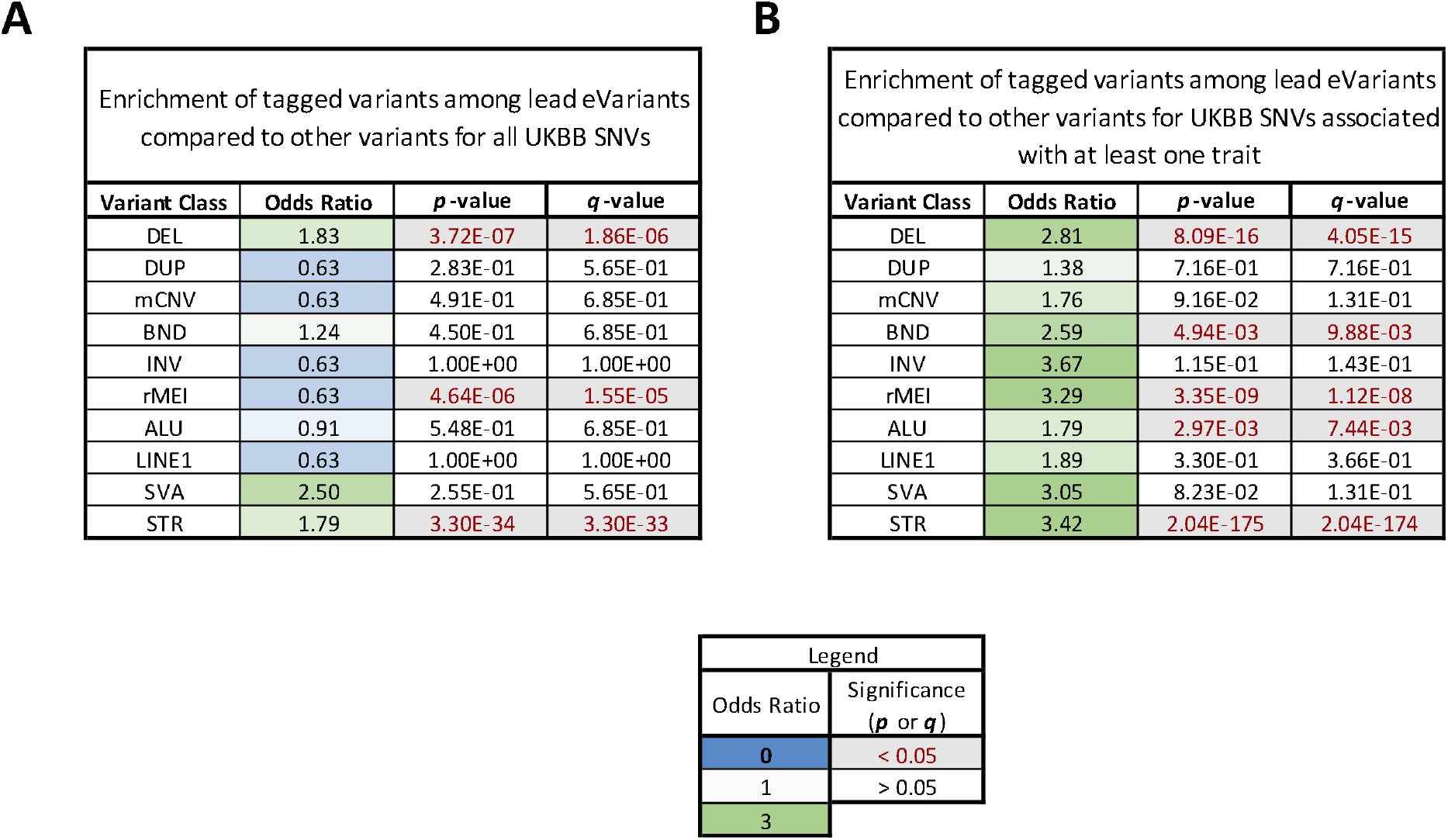
LD with UKBB variants and GWAS Traits. (A) Enrichment odds ratios, p-values and q-values for the likelihood of tested variants in the SV/STR-only eQTL analysis from each class to be in strong LD (R^2^>0.8) with a UKBB variant (± 25kb of SV/STR) (A) or a UKBB variant that is strongly associated with at least one GWAS trait (p-value < 5e-8) (B) when they are or are not lead eVariants.

## Supplementary Tables

**Table S1: Sample Information.** Information describing 398 RNA-seq samples included in variant calling including cell type, subject, study, family annotation, monozygotic twin status, and presence in the unrelated set of samples.

**Table S2. Joint-eQTL Mapping Results.** Lead associations including ties for all 11,197 eGenes discovered in the joint-eQTL analysis.

**Table S3. SV/STR-only eQTL Mapping Results.** Lead associations and all significant associations for 6,996 eGenes as well as annotation of the proximity of variants to promoter capture loop anchors (closest anchor-proximal or distal, orientation within or outside of loop, distance to anchor). These data were used in Figures 3-6.

**Table S4. LD with UKBB GWAS Traits.** Information about the i2QTL SV/STR calls that are in strong LD with a nearby UKBB variant that is significantly associated with at least one trait (p < 5e-8).

**Table S5:**
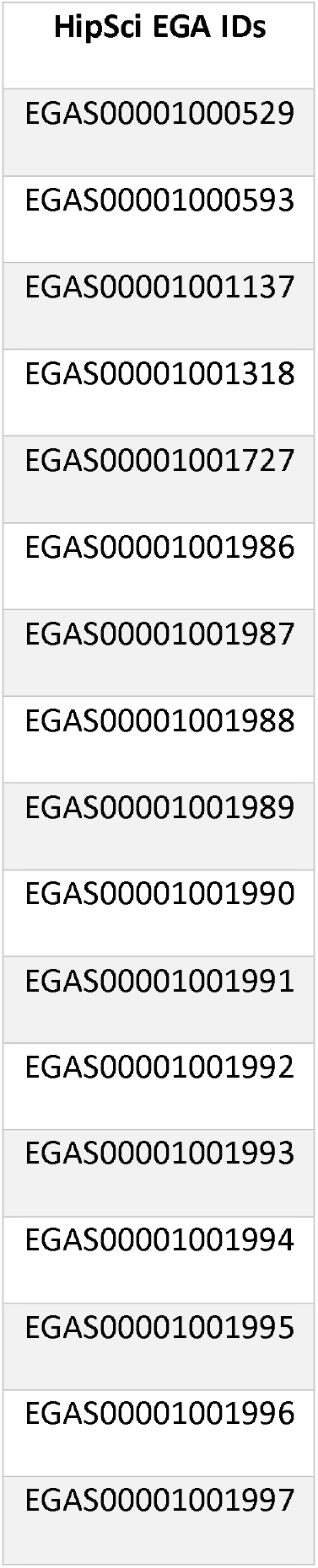
Hipsci EGA Projects. EGA IDs for HipSci data included in the i2QTL full sample set.

